# Designer indicators for two-photon recording of subthreshold voltage dynamics

**DOI:** 10.64898/2026.04.08.717226

**Authors:** M. Land, M. Galdamez, V. Villette, J. Zhu, X. Lu, M. Marosi, S. Yang, A. J. McDonald, X. Dong, E. Zaabout, H. Liu, Z. Liu, KL. Colbert, S. Lai, M. Shorey, A. Ayon, J. Bradley, C. Mailhes-Hamon, RG. Natan, J. Zhong, R. Kroeger, RG. Law, N. Hakam, CL. Smith, M. Hu, S. Tabb, B. Dudok, N. Ji, L. Bourdieu, J. Reimer, F. St-Pierre

## Abstract

Subthreshold voltage dynamics are critical for neuronal information integration, yet they remain understudied in vivo due to the limitations of current tools. While genetically encoded voltage indicators (GEVIs) offer a promising alternative, their application for deep-tissue recording using two-photon (2P) microscopy—a preferred method for deep-tissue recording—has been hindered by insufficient sensitivity for detecting millivolt-scale subthreshold signals. Here, we refined our multiparametric two-photon high-throughput screening platform to develop two novel GEVIs, JEDI3sub and JEDI3hyp, tailored explicitly for subthreshold voltage detection. Through fast 2P optical recording in awake, behaving mice, we demonstrated the superior sensitivity of JEDI3 indicators compared to JEDI-2P. We also showed that JEDI3sub can track population-level subthreshold optical tuning, while JEDI3hyp reliably captured subthreshold dynamics associated with sharp-wave ripple oscillations in hippocampal PV interneurons. Finally, JEDI3hyp facilitated extended imaging of brain-state-dependent, millivolt-scale subthreshold voltage changes across deep-layer somas, fine dendritic structures, and diverse cell types. By addressing the critical gap in 2P optical recording of subthreshold voltage dynamics, JEDI3 indicators open new avenues for studying neural information processing and its alterations in health and disease.

## Introduction

Neurons process and transmit information by regulating membrane voltage. Neurotransmitters at synapses can produce graded, subthreshold voltage changes that propagate through dendrites, where they are filtered, attenuated, or amplified. At the axon initial segment (AIS) near the cell body, neurons convert these signals into all-or-none voltage spikes called action potentials (APs). Monitoring subthreshold signals is essential to understand how neurons integrate inputs, detect functional synaptic connections, and explore how learning, disease, and aging affect synaptic strength^1–3^.

Although tools to monitor spiking activity are comparatively more developed, techniques to detect subthreshold events in genetically defined neurons deep within tissue remain limited. Whole-cell patch-clamp electrophysiology can record subthreshold events with high temporal resolution, but it is technically demanding in awake, behaving animals and unsuitable for small dendrites and axons. Furthermore, this technique often cannot unambiguously identify the recorded neuron’s cell type^4^. Genetically encoded calcium indicators (GECIs) enable cell-type-specific imaging of calcium dynamics underlying spiking activity across large brain volumes^5^. However, because GECIs detect calcium influx associated with APs, they generally cannot report subthreshold activity^6^.

Genetically encoded voltage indicators (GEVIs) offer a promising alternative, translating membrane potential changes into fluorescence signals^7^. We recently developed JEDI-2P, a GEVI capable of prolonged (>40 min) optical two-photon (2P) voltage recordings in deep tissue (layer 5) of awake, behaving animals^8^. However, its sensitivity to subthreshold voltage dynamics is limited, with fractional fluorescence responses of only ∼0.7%/mV between -70 and -50 mV^8^. Improved detection of millivolt-scale voltage changes is essential for studying phenomena such as inhibitory and excitatory postsynaptic potentials (IPSPs and EPSPs) and low-frequency oscillations linked to cortical synchronization and firing rates^9–15^. Two-photon optimized sensors with enhanced sensitivity to subthreshold depolarizations and hyperpolarizations are urgently needed to expand the GEVI toolbox suitable for in vivo physiology.

We present advancements in our two-photon high-throughput screening platform, enabling the evolution of GEVIs optimized for detecting subthreshold voltage dynamics. This system yielded two novel GEVIs, JEDI3sub and JEDI3hyp, each with enhanced responses to subthreshold voltages and distinct ranges of maximal membrane potential sensitivity. The JEDI3 sensors reliably captured subthreshold changes during spontaneous activity, stimulus-driven responses, and hippocampal short-wave ripples. Notably, JEDI3hyp tracked millivolt-scale, brain state-dependent somatic and dendritic subthreshold voltage changes in Layer-5 pyramidal neurons. Together, the JEDI3 indicators enabled robust monitoring of subthreshold voltage dynamics across various brain regions, cell types, and imaging modalities.

## Results

### Multiparametric high-throughput optimization of GEVIs for 2P recording of subthreshold voltage dynamics

To create an indicator with greater sensitivity to subthreshold voltage changes, we selected JEDI-2P as the base sensor, owing to its proven ability to perform sustained deep-tissue 2P spike recordings in vivo^8^. Our aim was to enhance JEDI-2P’s responsiveness to subthreshold voltages while preserving or improving other critical properties. We employed a 2P multiparametric high-throughput screening platform that measures sensor’s responses to brief (∼10-ms) voltage signals, brightness, and photostability^8^. We expressed sensor variants in HEK293-Kir2.1 cells, maintained at a resting membrane potential of -82.6 mV (±2.9 mV, 95% CI, n= 11) using an extracellular solution containing 5 mM potassium. This potential was chosen because mammalian neurons rarely hyperpolarize below -85 mV^16^. Cells were depolarized using electrical field stimulations of varying strengths, hypothesizing that selecting variants with pronounced responses to weak stimulations (Fig. S1A) would yield GEVIs optimized for detecting subthreshold voltage changes.

We screened nearly one hundred libraries arrayed in 96-well plates, including 79 targeting a single site by saturation mutagenesis and 17 combining promising mutations. Of the 79 mutated residues, 48 resided in the voltage-sensing domain, and 31 in the GFP moiety (Fig. S1B). The two best-performing indicators, JEDI3hyp and JEDI3sub, demonstrated larger responses to both partial and maximal field stimulations compared with JEDI-2P, while retaining comparable brightness and photostability (Fig. S1C-D).

JEDI3hyp and JEDI3sub differed from JEDI-2P by 7 and 8 mutations, respectively (Fig. 1A, Fig. S1B). Three mutations occurred in the voltage-sensing domain: S144V and M395T, located in the extracellular loop near the cpGFP insertion site, and R88S, positioned N-terminal to the S2 segment. Four mutations were in the GFP domain. Of these, N150R (GFP N146) lies near the chromophore and a GFP insertion point and is shared with the GEVI ASAP3^17^. K162R emerged from screens of glutamate indicator iGluSnFr mutations^18^. T209A (GFP T205) may influence excited-state proton transfer^19^. C295S (GFP C48) modifies a β-barrel cysteine, potentially preventing aberrant folding in the endoplasmic reticulum^20^. Lastly, the V399F mutation was identified in libraries aiming to shift JEDI3hyp’s sensitivity to more polarized potentials, aligned it more closely with the physiological range of neuronal voltages^21^. Residue 399 lies at the N-terminus of S4, a transmembrane segment critical for voltage sensing^22^.

**Figure 1.**
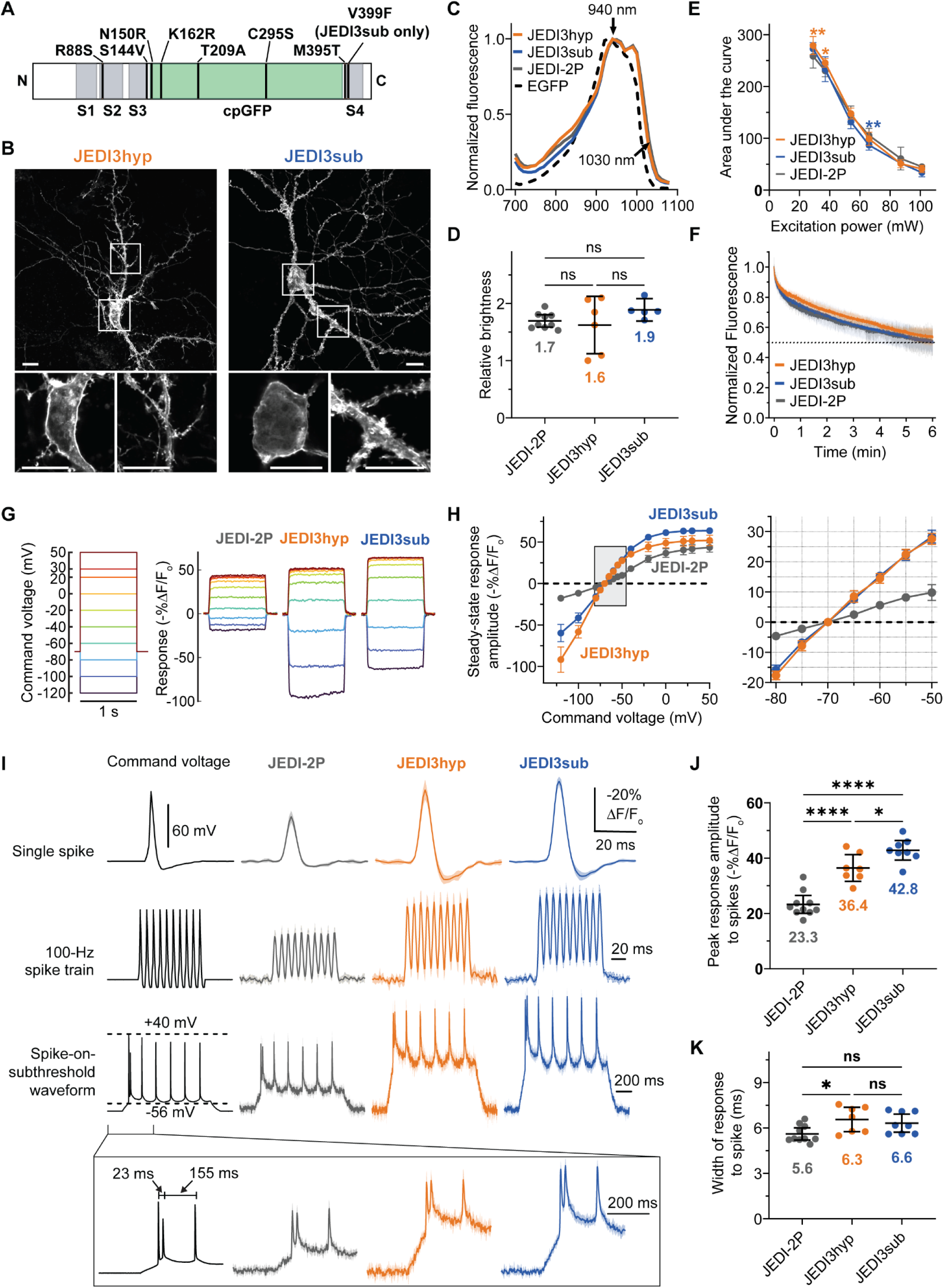
JEDI3 indicators exhibit enhanced sensitivity to subthreshold voltages while maintaining comparable brightness and photostability as JEDI-2P *in vitro*. (**A**) JEDI3hyp and JEDI3sub mutations relative to the parental JEDI-2P sequence. *Gray*, transmembrane segments of the voltage-sensing domain. *Green*, the circularly permuted GFP domain. (**B**) JEDI3 indicators efficiently targeted the plasma membrane of representative cultured rat E18 hippocampal neurons. Soma zoom-in images were taken from single confocal Z slices, while other images are confocal Z-stack maximal projections. Scale bars, 10 μm. (**C**) 2P excitation spectra in HEK293-Kir2.1 cells. GEVIs showed a peak excitation at 940 nm, red-shifted compared with EGFP (930 nm). Spectra were normalized to their respective peaks. Laser pulses were not pre-compensated for dispersion in the microscope optical path. Data are the mean of n = 14 fields of view per construct, each containing >100 HEK293-Kir2.1 cells. (**D**) JEDI3 indicators displayed comparable brightness to JEDI-2P at -70 mV. Sensor fluorescence was normalized to the brightness of the covalently linked mBeRFP. Black lines denote the mean brightness of n = 5-9 cells/indicator. *, *p*<0.05; ****, *p*<0.0001; n.s., *p*>0.05, one-way ANOVA with Tukey’s multiple comparisons test. (**E**) Representative photobleaching trace using 37-mW excitation light at the sample plane. Fluorescence was normalized to the intensity at t = 0. (**F**) The experiment in panel E was repeated for different excitation powers. The area under the curve (AUC) for each of the 6-min timecourses was quantified and normalized to the maximal AUC. Data is the mean from n = 3 independent transfections per GEVI in HEK293-Kir2.1 cells. Statistical comparisons between JEDI-2P and JEDI3hyp (*orange* *) and JEDI3sub (*blue* *): *, *p*<0.05; **, *p*<0.005; two-way ANOVA with Dunnett’s multiple comparisons test. (**G**) JEDI3 indicators produced larger steady-state responses to voltage steps than JEDI-2P, particularly for the subthreshold voltages. Mean responses are displayed with traces smoothed by a 47.6-ms moving average. n = 7-10 cells/variant. (**H**) Peak steady-state responses to step voltages. (**I**) JEDI3 indicators generated larger responses to spike waveforms than JEDI-2P. Traces are the mean responses. To mimic the properties of layer 2/3 cortical neurons at room temperature, single spikes and APs in trains had a 2-ms full width at half maximum^1^. The bottom-row waveform was adapted from a recording of mouse L5 pyramidal neurons and displays slightly wider spikes (3-4 ms). n = 7-10 HEK293A cells/variants. Imaging was acquired by resonant scanning at 440 Hz. (**J,K**) Fluorescence response amplitude (J) and width (K) to single spike waveforms. Statistics as in panel (D). For all panels, data was collected at 23°C with the laser set to 940 nm, and error bars or shaded areas denote the 95% CI.

### In vitro characterization of JEDI3hyp and JEDI3sub

To characterize JEDI3hyp and JEDI3sub in vitro, we first confirmed that the introduced mutations did not impair expression or membrane localization in neurons, a critical prerequisite for in vivo applications. Proper localization at the plasma membrane is essential, as intracellularly trapped sensors can fluoresce but will typically fail to respond to plasma membrane voltage changes, reducing overall signal amplitude. Confocal microscopy revealed robust plasma membrane localization in the soma and dendrites of dissociated rat hippocampal neurons for both JEDI3 variants (Fig. 1B).

Next, we determined the two-photon excitation spectra of JEDI3hyp and JEDI3sub to identify the optimal wavelength for further characterization. Both indicators exhibited excitation spectra similar to JEDI-2P, with a peak at 940 nm and a broad profile maintaining over 85% of peak excitation between 920 and 1,000 nm. While most in vivo applications were conducted at or near the peak wavelength, illumination at red-shifted wavelengths such as 980-1,000 would reduce scattering and could provide enhanced deep-tissue performance. For example, illumination at 980–1,000 nm can provide 90-95% of excitation at 940 nm. All JEDI indicators demonstrated significant excitation (25-32%) at 1,030 nm, enabling compatibility with fiber lasers used in emerging techniques such as scanless^23^, SLAP^24^, and FACED microscopy^25^.

We previously demonstrated that JEDI-2P possesses high brightness and photostability, supporting continuous in vivo recordings for over 30 minutes across various microscopy techniques^8^. JEDI3 sensors maintained comparable brightness (Fig. 1D) and photostability (Fig. 1E) to JEDI-2P, indicating their potential for similarly extended in vivo recordings. Fluorescence traces for all JEDI indicators revealed bi-exponential photobleaching kinetics, with an initial rapid bleaching phase followed by a slower phase (Fig. 1F).

We then assessed whether JEDI3 indicators displayed enhanced responses to subthreshold voltage changes, as predicted by our screens. Using two-photon voltage imaging combined with whole-cell voltage clamp, we found that JEDI3 indicators produced 2.7 and 3.6 times larger responses than JEDI-2P to 10-mV depolarizations and hyperpolarizations, respectively (Fig. 1G-H). Responses to 100-mV voltage steps from -70 mV were also significantly larger: JEDI3hyp and JEDI3sub produced responses of 51.5% and 63.5%, respectively, compared with 41.8% for JEDI-2P.

We quantified indicator kinetics to assess how effective GEVI fluorescence tracks voltage dynamics. Fast depolarization kinetics are particularly critical for accurately capturing spikes. For voltage steps corresponding to suprathreshold events (-70 or -45 mV to +30 mV), JEDI3hyp and JEDI3sub maintained the short response time constants of JEDI-2P (Table 1). For subthreshold (-70 to -45 mV) and hyperpolarization (-70 to -100 mV) steps, JEDI3hyp and JEDI3sub exhibited slightly slower kinetics than JEDI-2P. Notably, the kinetic rates of indicators derived from identical or orthologous voltage-sensing domains exhibit an increase with rising temperature^8,26^. Thus, indicators are expected to respond faster at physiological temperatures in mice than at 33°C, the temperature used here due to challenges in maintaining robust patch seals at higher temperatures.

**Table 1.** Indicator kinetics at 33°C. Values are means ± 95% CI. For voltage steps from -70 to +30 mV, and +30 to -70 mV: n = 7 (JEDI-2P), 15 (JEDI3hyp) and 10 (JEDI3sub) cells. For other voltage steps: n = 9 (JEDI-2P), 8 cells (JEDI3hyp) and 10 (JEDI3sub) cells. HEK293A cells were used.

|  | JEDI-2P | JEDI3hyp | JEDI3sub |
| --- | --- | --- | --- |
| **-70 to +30 mV** |  |  |  |
| Tau fast (ms) | 0.55 ± 0.06 | 0.48 ± 0.05 | 0.63 ± 0.05 |
| Tau slow (ms) | 7.57 ± 3.59 | 7.97 ± 4.29 | 5.31 ± 1.39 |
| Fast component (%) | 85.1 ± 3.5 | 90.3 ± 3.7 | 90.8 ± 1.4 |
| **+30 to -70 mV** |  |  |  |
| Tau (ms) | 1.20 ± 0.11 | 2.11 ± 0.21 | 2.23 ± 0.19 |
| **-45 to +30 mV** |  |  |  |
| Tau (ms) | 1.36 ± 0.15 | 1.69 ± 0.45 | 1.18 ± 0.15 |
| **+30 to -45 mV** |  |  |  |
| Tau (ms) | 1.68 ± 0.12 | 2.90 ± 1.52 | 2.92 ± 0.22 |
| **-70 to -45 mV** |  |  |  |
| Tau (ms) | 1.04 ± 0.06 | 2.59 ± 0.39 | 2.25 ± 0.17 |
| **-45 to -70 mV** |  |  |  |
| Tau (ms) | 0.96 ± 0.18 | 3.00 ± 0.35 | 2.45 ± 0.14 |
| **-100 to -70 mV** |  |  |  |
| Tau (ms) | 1.46 ± 0.23 | 3.14 ± 0.35 | 2.53 ± 0.17 |
| **-70 to -100 mV** |  |  |  |
| Tau (ms) | 1.04 ± 0.24 | 3.43 ± 0.26 | 2.27 ± 0.10 |

Finally, we assessed how JEDI3 kinetics and steady-state responses influenced spike and subthreshold detectability. JEDI3hyp and JEDI3sub produced peak responses to 2-ms full-width at half-maximum (FWHM) AP waveforms that were 56% and 84% larger, respectively, than JEDI-2P (Fig. 1I, first row; Fig. 1J). JEDI3hyp produced slightly longer responses to 2-ms FWHM AP waveforms when compared with JEDI3sub and JEDI-2P (Fig. 1K). However, all JEDIs reliably tracked 100-Hz spike train waveforms (Fig. 1I, second row), consistent with their overall rapid kinetics. When tested with spike-on-subthreshold waveforms recorded from mouse L5 pyramidal neurons, JEDI3 sensors exhibited markedly enhanced responses to subthreshold components compared with JEDI-2P (Fig. 1I, third row).

### JEDI3 sensors exhibit greater subthreshold voltage responses than JEDI-2P in awake behaving mice

We evaluated whether the improved subthreshold voltage dynamics of JEDI3 indicators observed in vitro extended to in vivo preparations. Benchmarking was performed using ultrafast local volume excitation (ULoVE) microscopy, a 2P technique based on acousto-optic deflectors that allows high (>7.1 kHz) temporal resolution and sensitivity for single-cell recordings^17^. We recorded layer 2/3 cortical neurons expressing GEVIs at depths of 69.2–304 µm (mean ± SD: 178 ± 64.4 µm; n= 42 cells from 8 mice) in head-fixed, behaving mice (Fig. 2A, B). All indicators were appended with the Kv motif, corresponding to the PRC domain from the potassium channel Kv2.1, to preferentially localize indicators at the soma^27^. Suprathreshold (AP) response amplitudes were derived from spike-triggered averages (Fig. 2C), while subthreshold responses were quantified as the range of the low-pass-filtered trace.

**Figure 2.**
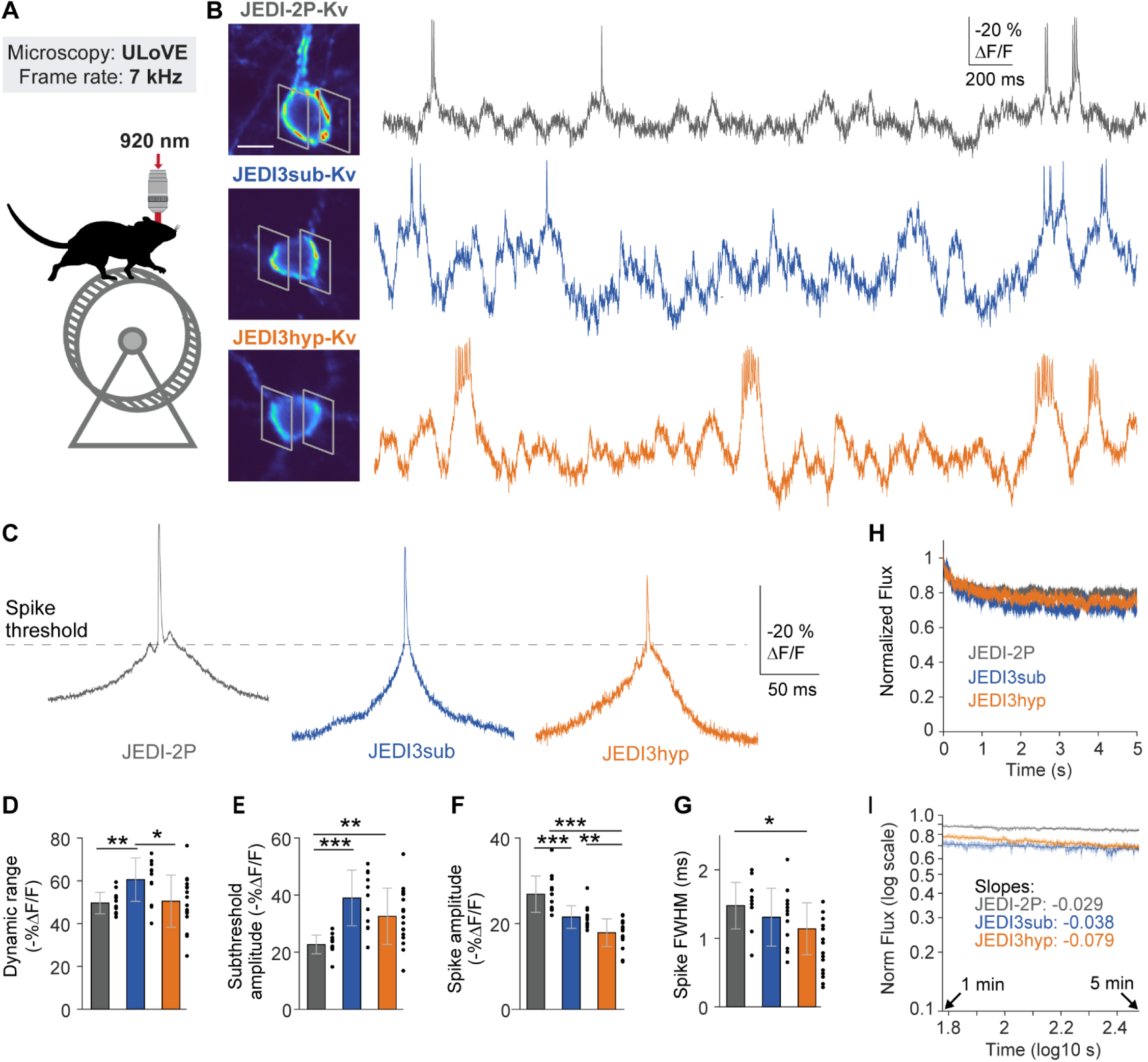
JEDI3 indicators report subthreshold dynamics with greater amplitude than JEDI-2P in cortical neurons of awake, behaving mice. (**A**) Experimental design schematics. Mice were head-fixed and allowed to move on a self-propelled treadmill while 2P-induced fluorescence was recorded using ULoVE microscopy. (**B**) Representative responses from layer-2/3 neurons transduced using AAVs with Cre-dependent GEVI expression. Soma-restricted expression was achieved using the Kv motif. The two gray polygons in each image (left) represent the approximate boundaries of the ULoVE excitation patterns. Scale bar,10 µm. (**C**) Mean optical spike waveforms computed from isolated spikes for the recordings in panel (B). (**D-G**) Quantitative GEVI benchmarking. n = 13 (JEDI-2P), 12 (JEDI3sub) and 17 (JEDI3hyp). *, p<0.05; **, p<0.01; and ***, 0.001, two-sample Wilcoxon rank sum test. Error bars represent the SD. (**H, I**) Photostability of JEDI3 indicators was comparable to JEDI-2P, as evidenced by similar changes in average photon flux during the initial fast bleaching phase (H) and the remaining fluorescence trace (I). Sample sizes as in panels (D-G).

Among the indicators, JEDI3sub demonstrated the highest overall performance, with a dynamic range (subthreshold + spike) of 60.6%, 1.2-fold greater than that of JEDI-2P (49.6%, Fig. 2D). JEDI3sub’s subthreshold responses were 1.7-fold larger than JEDI-2P’s (Fig. 2E). However, unlike in vitro, JEDI3sub’s spike responses were 0.8-fold those of JEDI-2P, with a 21.6% ΔF/F_0_ response amplitude compared with JEDI-2P’s 26.9% (Fig. 2F). JEDI3hyp also exhibited enhanced subthreshold responses compared with JEDI-2P, but both its subthreshold and spike responses were lower than JEDI3sub’s. Spike widths were similar across all indicators, with JEDI3hyp producing slightly narrower spikes than JEDI-2P (Fig 1G), while the opposite trend was observed *in vitro* (Fig. 1K). Differences in GEVIs’ relative spike response amplitudes between our *in vitro* and *in vivo* datasets could arise from variation in multiple factors including temperature (Fig. S2), spike waveforms, and acquisition methods.

Photobleaching was comparable across indicators, except for a slightly larger fast photobleaching component in JEDI3 sensors during the first few seconds of illumination (Fig. 2H). This was followed by a long, slow photobleaching component, characterized by the slope of log-log plotted data (Fig. 2I). Although JEDI3hyp exhibited a marginally larger photobleaching slope compared to JEDI3sub and JEDI-2P, this difference likely reflects variations in expression and in optical recording depth (JEDI2P: 136±35 µm, JEDI3hyp: 203±67 µm, JEDI3sub :188±66 µm) and thus illumination intensity *in vivo*.

A recently reported indicator, ASAP5, demonstrated the ability to detect single-trial mEPSPs in cultured neurons under one-photon illumination^28^. We compared the performance of JEDI3 indicators to ASAP5 using ULoVE in vivo, using equivalent experimental conditions, including brain structure, behavioral task, microscopy, and acquisition parameters. To avoid bias from potential differences in expression levels due to variations in viral capsid types or production methods, we focused exclusively on fractional fluorescence responses. Both JEDI3sub and JEDI3hyp outperformed ASAP5 in subthreshold response amplitudes, achieving 1.8-fold and 1.5-fold improvements, respectively (JEDI3sub: 39.0 +/- 9.7, JEDI3hyp: 32.5 +/- 9.9, ASAP5: 22.8 +/- 2.9 %ΔF/F_0_). JEDI3sub displayed statistically identical spike responses as ASAP5, while JEDI3hyp produced responses that were 0.79-fold smaller (JEDI3sub: 21.6 +/- 2.6, JEDI3hyp: 17.9 +/- 3.2, ASAP5: 22.8 +/- 2.9 %ΔF/F_0_). Spike widths for all indicators were statistically indistinguishable.

### Population-level subthreshold voltage imaging reveals correlated neural activity

We next sought to evaluate JEDI3 indicators to record population-level subthreshold voltage dynamics. We expressed the soma-targeted JEDI3sub-Kv indicator in the visual cortex of awake mice (Fig. 3A), producing densely-labeled populations of neurons with excellent indicator expression at the plasma membrane (Fig. 3B). We leveraged Free-space Angular-Chirp-Enhanced Delay (FACED) 2P microscopy, which employs megahertz line scanning to enable the capture of voltage activity across large populations of neurons within the same focal plane^25,29^. Optical imaging was conducted at 400 or 769 Hz over a 320x400 µm field-of-view while mice were presented with drifting grating visual stimuli. JEDI3sub exhibited minimal photobleaching during ∼200-second imaging sessions (Fig. S3B).

**Figure 3:**
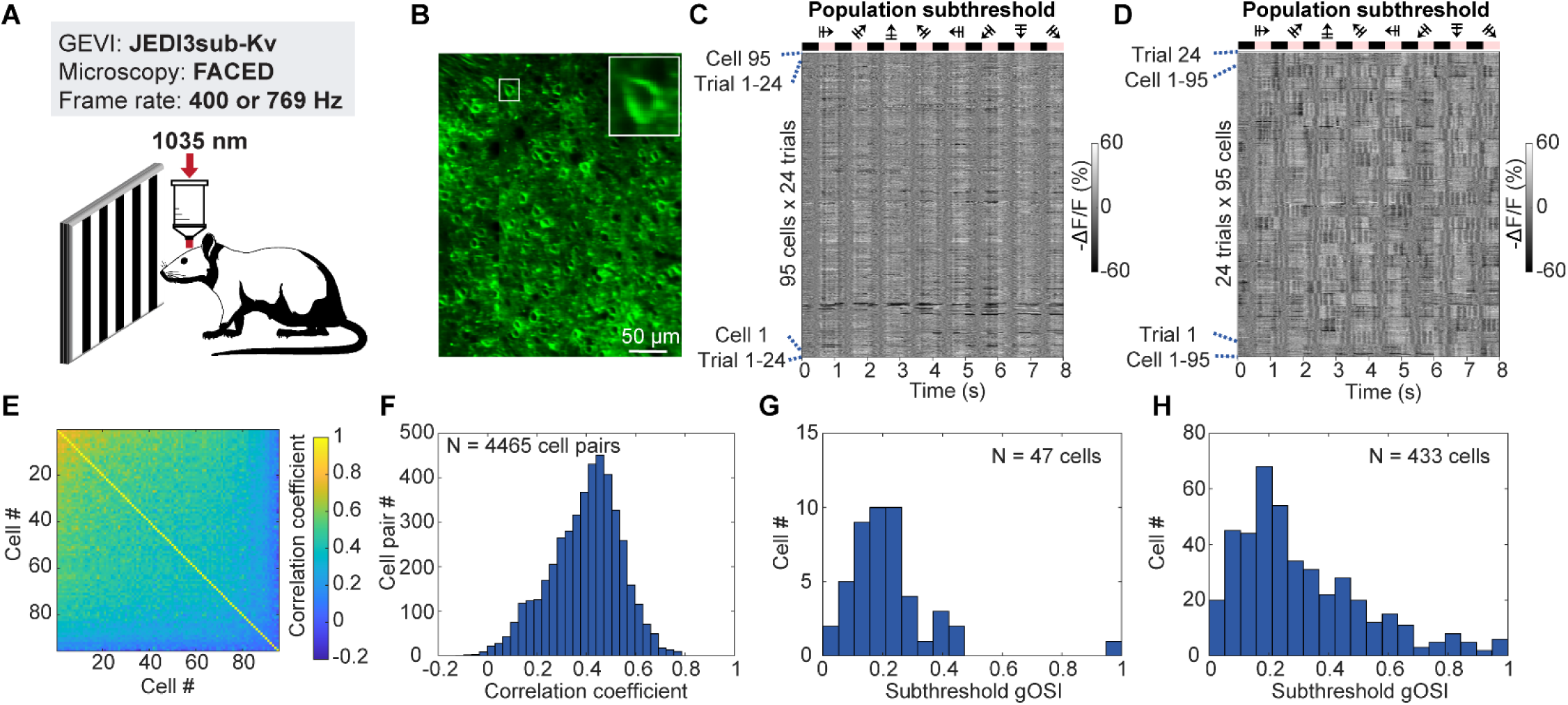
JEDI3sub tracks subthreshold optical tuning in 95 neurons in awake, behaving mice. (**A**) Experimental design schematics. AAVs expressing the soma-restricted indicator JEDIsub-Kv from the pan-neuronal promoter hSyn were injected into the visual cortex. FACED microscopy was used to record 2P fluorescence responses to drifting gratings presented to the right eye of head-fixed, awake behaving mice. The post-objective excitation power was set to 156 mW. (**B**) Representative 320 µm × 400 µm voltage imaging field of view showing JEDI3sub-Kv-expressing neurons. (**C, D**) Single-trial subthreshold traces from 95 cells and 24 trials, grouped by (**C**) cells and (**D**) trial. (**E, F**) Correlation analysis of subthreshold responses, with (**E**) displaying the Pearson’s correlation coefficient (CC) matrix for subthreshold responses between 4,465 cell pairs, with cells sorted by the mean CC and (**F**) a histogram showing the distribution of CC values. (**G, H**) Neuron orientation selectivity analysis. Histograms of subthreshold global orientation-selectivity indices (gOSI) for (**G**) n = 47 orientation-selective (OS) neurons from one mouse (example shown in panel B) and (**H**) 433 OS neurons across 13 fields of view from 2 animals.

**Figure 4.**
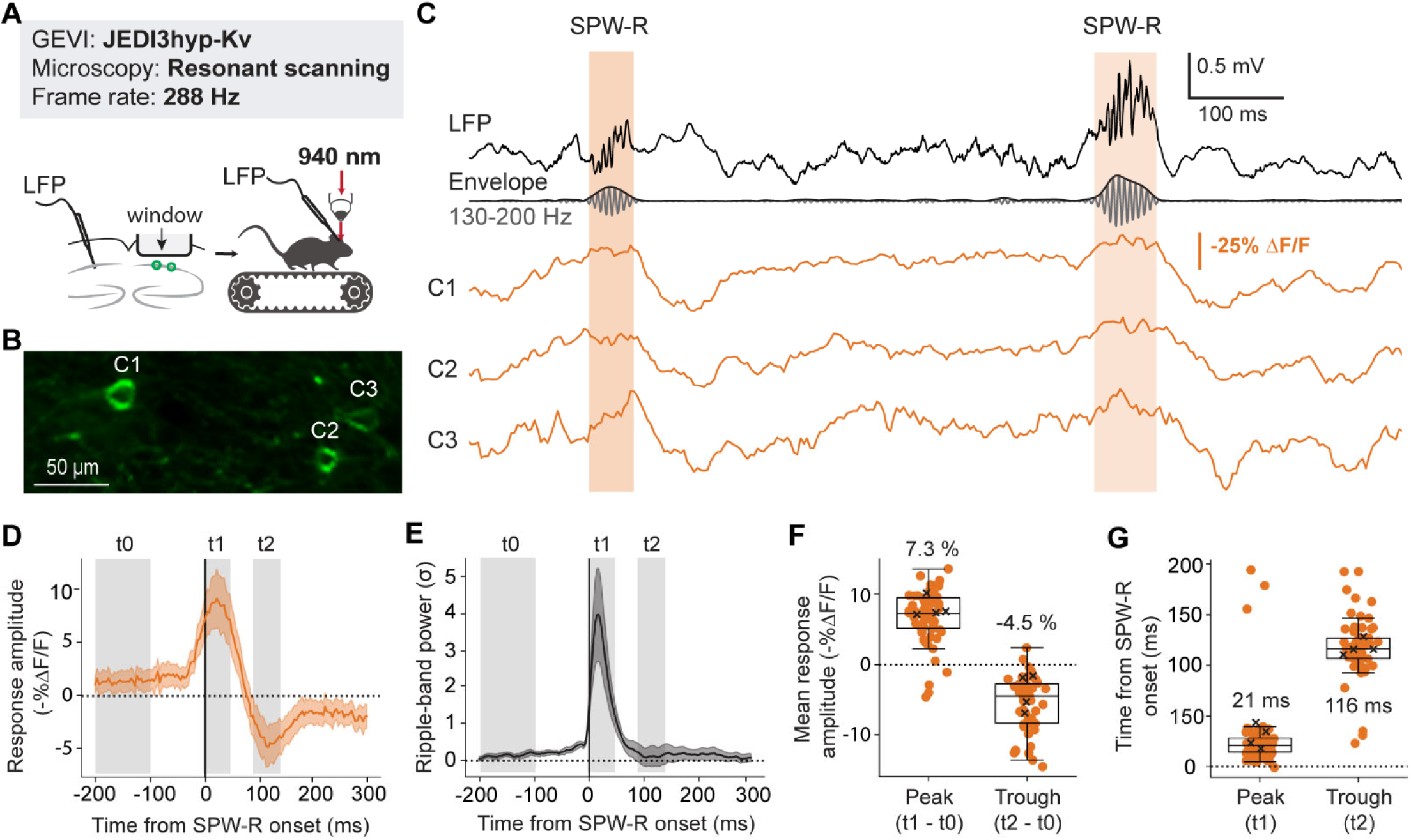
JEDI3hyp reveals subthreshold potential changes associated with spontaneous network oscillations *in vivo*. (**A**) Experimental design schematics. PV-Cre mice (n=4 females, P53-67) were injected with AAVs expressing Cre-dependent soma-restricted JEDI3hyp-Kv in PV interneurons (green circles). Mice were head-fixed to a self-propelled treadmill, allowing the mice to run or rest freely. 2P resonant scanning microscopy of the hippocampal CA1 region was conducted at a frame rate of 288 Hz using a laser set to 940 nm. Local field potentials (LFP) were recorded simultaneously with a bipolar electrode. (**B**) An example field of view in the CA1 pyramidal layer, representative of 24 regions imaged. Note the soma-restricted JEDI3hyp-Kv expression. (**C**) Fluorescence and LFP traces during sharp wave-ripples (SPW-Rs). Segments of the fluorescence traces from the three representative cells shown in (B), overlaid with the simultaneously recorded LFP. SPW-R events (orange bars) were detected as peaks in the envelope (black) of the ripple-band filtered trace (dark gray, 130-200 Hz). Fluorescence responses from the three cells from panel B (orange traces) show transient depolarizations during SPW-Rs and hyperpolarizations afterward. Fluorescence traces were processed for visualization using an exponentially weighted moving average with a 25-ms decay. Note that the 288-Hz frame rate supports a larger field of view but does not resolve individual spikes or ripple oscillations. (**D,E**) Event-triggered PV+ interneuron fluorescence (D) and LFP ripple-band power (E) during SPW-Rs. Data is mean ± SEM. The baseline (t0), peak (t1, 21 ± 25 ms), and trough (t2, 114 ± 25 ms) are marked with gray bars and used to calculate the responses in (F,G). n = 60 cells and 4 mice. (**F,G**) Mean fluorescence response (F) and timing from SPW-R onset (G) of the peak and through in individual neurons (orange circles). Boxes show median ± interquartile range; whiskers show the nonoutlier range. Black X’s show averages by animal. Mean fluorescence increased during peaks (p = 0.00075) and decreased during troughs (p = 0.029; one-sided, one-sample t-test, n = 4 animals).

From 95 analyzed cells, >92% (88) displayed single-trial visually evoked subthreshold activity (Fig. 3C-D). This is substantially higher than the 49% of L2/3 neurons with drifting-grating-evoked suprathreshold activity measured by calcium imaging^30^, which does not report subthreshold responses. Grouping trials revealed significant biological variability, with each trial showing distinct patterns of synchronized population activity, likely related to brain state (Fig. 3D; Fig. S3C; see section regarding Fig. 5). Pairwise correlation analysis yielded a mean correlation coefficient of 0.40 across 4465 cell pairs (Fig. 3E-F). Similar findings were observed in a total of 1,230 neurons across 13 FOVs from 2 mice (two more example FOVs shown in Fig. S3D). Among neurons with visually evoked subthreshold activity, 53.4% exhibited orientation selectivity (Fig. 3G). Subthreshold global orientation-selectivity indices (gOSIs) displayed the population-level orientation selectivity distribution for the example FOV (Fig. 3G) and for 433 neurons from 2 mice (Fig. 3H).

**Figure 5.**
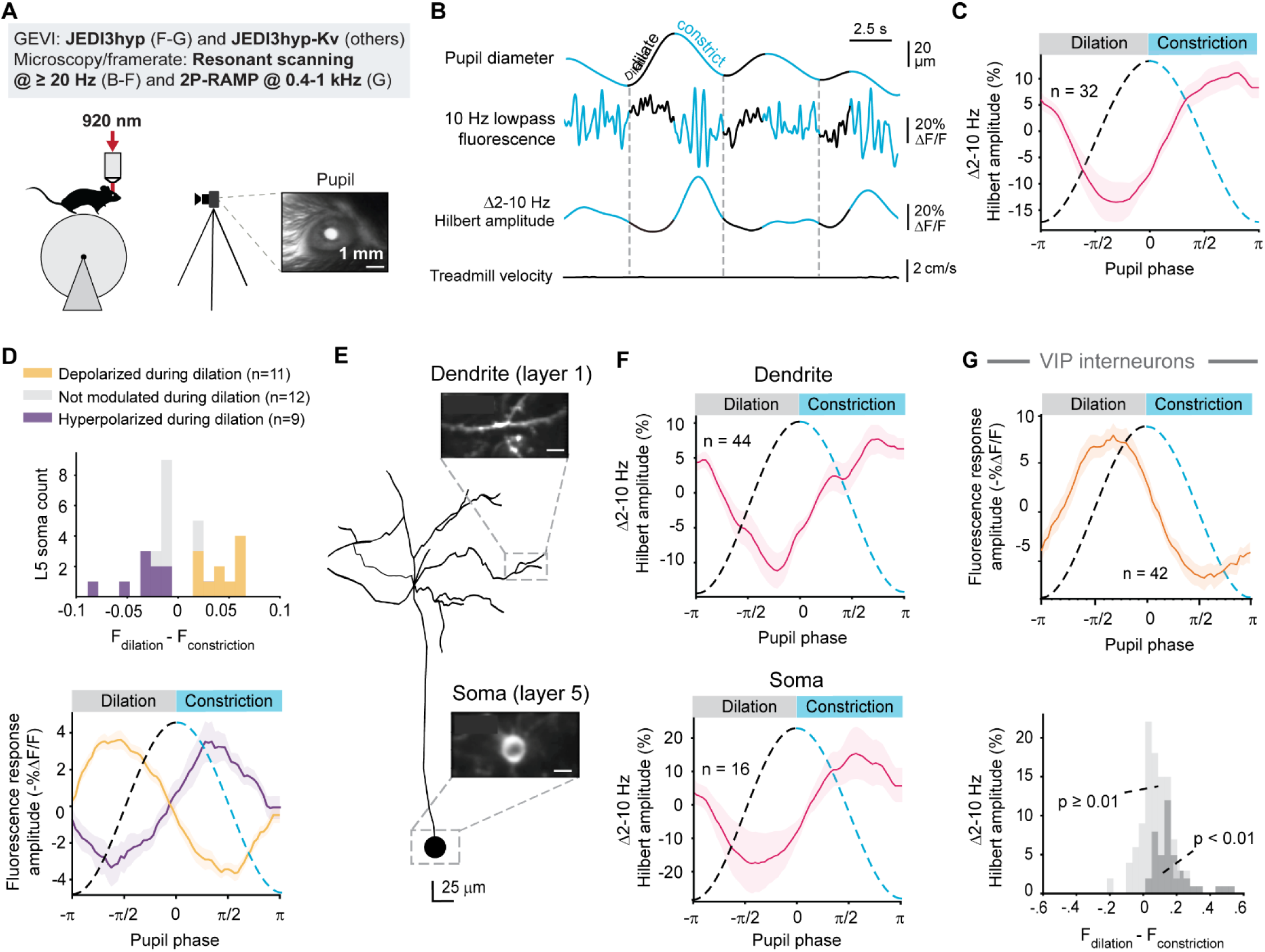
JEDI3hyp reliably reports voltage changes associated with brain state in different cell types, subcellular locations, and cortical layers. (**A**) Experimental setup schematics. AAVs expressing Cre-dependent JEDI3hyp or JEDI3hyp-Kv were injected into mice’s visual cortex. Mice were head-fixed and freely allowed to move on a cylindrical treadmill. Experiments were conducted using 2P resonant scanning (B-F) or random-access (G) microscopy using lasers set to 920 nm. Pupil diameters were simultaneously recorded. (**B**) Example layer-5 intratelencephalic pyramidal neuron (L5 IT-PN) showing how changes in pupil diameter (*top*) are associated with changes in 2-10 Hz voltage oscillations, as reported by JEDI3hyp (2^nd^ and 3^rd^ traces). Dilation and constriction periods are colored black and blue, respectively. Vertical dashed lines separate adjacent dilation/constriction cycles. This example shows a period of quiet wakefulness, without any treadmill movement (bottom, black). (**C**) Low-frequency activity in L5 IT-PNs (red trace, mean ± SEM) decreases following pupil dilation onset and peaks during constriction. n = 32 soma from four mice. (**D**) L5 IT-PNs can be subdivided into two functional subclasses exhibiting either hyperpolarization (purple, n=9) or depolarizations (yellow, n=11 soma) during pupil dilation (black). p < 0.05, Kruskal-Wallis test. (**E**) Example reconstruction of a JEDI3hyp-expressing L5 IT-PN neuron. An apical dendrite and its corresponding soma are shown. Scale bar, 20 µm. (**F**) The reduction of low-frequency membrane potential oscillations during pupil dilation is visible both in dendrites (*top*, n=44) and somas (*bottom*, n=16). Multi-depth imaging was achieved using a 2P-RAM mesoscope. (**G**) JEDI3hyp showed that VIP interneurons depolarize during pupil dilation, consistent with whole-cell patching studies (Reimer et. al., 2014). n = 111 total neural recordings from five mice. Of these, n = 42 (dark gray bars, bottom panel) showed the expected depolarization during dilation compared to constriction (p < 0.01, Kruskal-Wallis Test).

These results demonstrate JEDI3sub’s capacity to track subthreshold activity across large neuronal populations, enabling investigation into population-level information processing.

### JEDI3hyp tracks subthreshold dynamics associated with sharp-wave ripples in hippocampal interneurons

We investigated whether JEDI3 variants can capture behaviorally relevant subthreshold signals using conventional 2P resonant scanning microscopes. While these instruments offer limited fields of view at the high speeds required for voltage imaging, they remain widely used in neuroscience research. Validating JEDI3 indicators under these conditions is essential to assess their practicality and ensure compatibility with the imaging systems available in many neuroimaging labs.

First, we recorded Sharp-wave ripples (SPW-R), brief synchronous population events involved in cell sequence replay and which produce a stereotypical local field potential waveform in the CA1 hippocampal region^31,32^. In awake mammals, SPW-Rs are accompanied by a marked increase in the firing rates of parvalbumin-expressing (PV) interneurons that is phase-locked to the ripple oscillation^33^. PV interneuron spiking activity during SPW-R has been extensively studied using juxtacellular recordings and calcium imaging^34–36^. However, these methods do not report subthreshold voltage changes. In vivo patch-clamp has been used to characterize subthreshold responses in pyramidal cells^37^, but is difficult to perform on less abundant cell types such as interneurons. As a result, PV interneuron subthreshold activity related to SPW-Rs remains largely unexplored. Monitoring PV subthreshold potential changes may illuminate the activation of inhibitory inputs, which play an important role in SPW-R generation and termination^38,39^.

While both JEDI3 variants were expected to perform similarly in this context, we chose JEDI3hyp due to its potentially enhanced sensitivity to hyperpolarization (Fig. 1H). JEDI3hyp was transduced into the CA1 hippocampal region, expressed in PV interneurons using a PV-Cre mouse line, and imaged using a commercially available 2P resonant scanning microscope in awake behaving mice (Fig. 4A-B, S4). We set the acquisition frequency to 288 Hz, which while insufficient to capture spikes, enables the capture of graded membrane potential changes across a larger, 250x90 µm (256x48 pixel) field of view. JEDI3hyp tracked SPW-R-induced depolarizations and the following hyperpolarization of individual ripple-associated events in single neurons (Fig. 4C).

Next, we assessed whether the observed pattern of depolarization during SPW-Rs followed by hyperpolarization is a consistent feature across events, cells, and animals. Averaging the responses of all recorded PV interneurons, we found a clear depolarization during SPW-Rs (Fig. 4D), consistent with the previously reported increase in PV interneuron firing rates^33,34^. Peak depolarization occurred 21 ms after SPW-R onset, closely following the peak in ripple-band power at 17 ms (Fig. 4E-G). The average response of PV interneurons exhibited a post-SPW-R hyperpolarization, reaching a trough 116 ms after SPW-R onset. This post-SPW-R hyperpolarization mirrors a similar phenomenon reported in pyramidal cells, likely attributable to feedforward inhibition^37,40^.

The SPW-R-associated depolarization and subsequent hyperpolarization have also been observed using Voltron, a one-photon-compatible voltage indicator combining a fluorescent dye and a voltage-sensitive protein^41^, corroborating our findings. Our results underscore the capability of JEDI3hyp to accurately report behaviorally significant subthreshold signals in vivo under 2P microscopy and leveraging a fully genetically encoded indicator.

### JEDI3hyp reports brain state-dependent somatic and dendritic subthreshold voltage dynamics across cell types and cortical layers

We investigated whether JEDI3 indicators could report subtle subthreshold voltage changes in dendrites and deep cortical layers. In quiet awake animals, small but consistent subthreshold voltage fluctuations in cortical neurons correlate with pupil dilations and constrictions^9,16,42,43^. These dynamics, which are thought to influence state-dependent sensory processing and behavior, remain poorly understood. The current standard measurement technique, whole-cell patch clamping, is technically demanding, has low throughput, and is challenging in subcellular compartments and deeper cortical layers. We evaluated whether JEDI3 indicators could overcome these limitations by enabling rapid, reproducible optical measurements of voltage dynamics across cortical layers, cell types, and subcellular compartments.

Patch-clamp studies during quiet wakefulness have shown that pupil dilation reduces 2-10-Hz voltage oscillations in layer-2/3 pyramidal neurons, while constriction enhances them^9^. Using 2P voltage imaging with resonant-scanning microscopy, we confirmed that JEDI3hyp-Kv replicated these electrophysiological findings (Fig. 5A, S5A,B) despite voltage changes being less than 2 mV on average^9^. Critically, JEDI3hyp-Kv demonstrated sufficient sensitivity to detect subthreshold voltage dynamics in layer-5 intratelencephalic pyramidal neurons (IT-PNs), a population challenging to access with patch clamping due to their depth. Similar to layer-2/3 neurons, layer-5 IT-PNs exhibited 2-10 Hz oscillations with lower Hilbert amplitude during pupil dilation and increased amplitude during pupil constriction (Fig. 5B-C). Approximately 63% of layer 5 IT-PNs displayed significant baseline fluorescence differences between pupil dilation and constriction. Of these, about half hyperpolarized, while the others depolarized during dilation (Fig. 5D, S5C), suggesting two functional subclasses of neurons with differing responses to neuromodulators like acetylcholine and norepinephrine, which are strongly modulated during pupil dilation^44^.

*In vivo* patch clamping of dendrites is technically challenging, laborious, limited to larger-diameter dendrites, and typically restricted to one location per cell. We tested JEDI3hyp’s ability to overcome these limitations and visualize physiologically relevant voltage fluctuations in the apical dendrites of layer-5 neurons. Using a dual-recombinase approach^45^ for sparse JEDI3hyp expression, we matched layer 5 somas to their corresponding layer-1 dendrites. Because this application required recording away from the cell body, we used JEDI3hyp without the Kv soma-restriction motif.

We conducted simultaneous multi-plane resonant scanning recordings from layer-5 somas and 1-6 areas with their corresponding layer-1 apical dendrites for over 30 minutes of continuous illumination (Fig. S5D). JEDI3hyp maintained photostability across imaging depths and laser intensities (Fig. S5E). These recordings revealed comparable low-frequency subthreshold voltage dynamics during pupil dilation and constriction in dendrites and soma (Fig. 5E-F). These results demonstrated JEDI3hyp’s utility in visualizing brain-state-dependent dendritic signals over extended periods.

Genetically encoded voltage indicators like JEDI3hyp also facilitate cell type-specific studies. We deployed JEDI3hyp to examine voltage changes in two GABAergic interneuron subtypes: vasoactive intestinal peptide-expressing (VIP) and somatostatin-expressing (SOM) cells. VIP interneurons depolarized during pupil dilation and hyperpolarized during constriction, consistent with patch-clamp findings (Fig. 5G, S5F)^9^. SOM cells showed more variable responses, reflecting their functional heterogeneity reported in electrophysiological studies (Fig. S5F)^9^. These results demonstrate JEDI3hyp’s sensitivity, given that maximal pupil-associated voltage changes in VIP and SOM cells have been measured at 2–3 mV on average^9^. More broadly, these results also establish JEDI3hyp as a robust tool for extended 2P imaging of brain-state-dependent subthreshold voltage changes across deep-layer somas, fine dendritic structures, and diverse cell types.

## Discussion

In this study, we introduced JEDI3sub and JEDI3hyp, 2P-optimized GEVIs for enhanced subthreshold voltage detection. By addressing a critical gap in the 2P imaging toolbox, which has historically emphasized spike detection, these sensors enable a deeper understanding of neuronal dynamics across various spatial and temporal scales.

### Advancement in biosensor engineering

The enhanced subthreshold detection capabilities of JEDI3 sensors were achieved through a deliberate and iterative high-throughput protein engineering strategy. We modified our 2P screening protocol to include field stimulation pulses with varying strengths and durations. This refinement enabled the prioritization of indicators with robust responses to weaker and shorter stimulations, which are representative of smaller voltage changes. This methodological innovation lays the groundwork for developing voltage indicators tailored to distinct physiological voltage ranges.

The mutations introduced into JEDI3 sensors often impacted multiple performance characteristics simultaneously, revealing critical trade-offs between key metrics (Fig. S1D). For instance, the S3-S4 loop mutations S144V and M395T, were pivotal in enhancing subthreshold detection but resulted in reduced brightness at resting membrane potential, likely due to a left-shifted fluorescence response curve. The perichromophore mutation N150R was introduced to restore brightness, although it also diminished responsiveness to partial field stimulations. These findings highlight the importance of balancing and simultaneously screening for multiple performance metrics during indicator optimization.

### Impact on neuroscience

JEDI3 sensors demonstrated utility in detecting subthreshold voltage dynamics across different brain areas, cell types, and subcellular compartments. Their use in advanced imaging methods enabled the simultaneous monitoring of subthreshold activity in approximately one hundred cells (Fig. 3), significantly enhancing the scale of optical recordings. Even with conventional resonant-scanning microscopy, JEDI3 sensors outperformed whole-cell patch-clamping by enabling the rapid and simple sequential recording of groups of neurons.

Our experiments revealed previously uncharacterized functional subclasses within layer-5 IT-PNs interneurons, likely driven by differential responses to cortical neuromodulators such as acetylcholine and norepinephrine. These findings align with prior studies linking neuromodulatory transients to pupil dilation and the biphasic responses of layer-5 pyramidal neurons to cholinergic input^44,46–48^. These results demonstrate the value of JEDI3 sensors to uncover novel neuronal dynamics, opening new avenues for understanding the roles of subthreshold activity in information processing.

### Potential for broader adoption of JEDI3 indicators

Subthreshold voltage dynamics occur at slower timescales than action potentials, allowing existing resonant scanning imaging systems, often used for calcium imaging, to be repurposed for JEDI3-based voltage imaging. Emerging optical recording technologies capable of ultra-high temporal resolution (Fig. 2), large-population imaging (Fig. 3), and multi-plane recordings (Fig. 5E,F), further expand the potential applications of JEDI3 sensors. For example, these tools make it possible to investigate the interactions between subthreshold dynamics and sensory or behavioral inputs across multiple layers and cell types simultaneously.

### How to choose between JEDI3sub and JEDI3hyp

Benchmarking experiments highlighted distinct advantages for each sensor (Fig.1, 2). JEDI3sub is recommended in most situations, given its superior responses to subthresholds voltage changes and robust spike detection. JEDI3hyp exhibited enhanced subthreshold sensitivity and exceptional performance at voltages below -80 mV, particularly at elevated temperatures (≥33°C), where its sensitivity advantage over JEDI3sub is most pronounced (Fig. S2). However, this improvement came at the cost of significantly diminished spike responses (Fig. 2C). focusing exclusively on subthreshold responses and involving hyperpolarized potentials below -80 mV. For instance, it may be particularly useful for studying the voltage dynamics of astrocytes, which can exhibit resting membrane potentials as low as -90 mV^49^.

### Limitations, future directions, and broader implications

While JEDI3 indicators have significantly enhanced the sensitivity of subthreshold voltage detection, further advancements in response amplitude and brightness are necessary to enable robust voltage recordings across larger populations of somata, smaller subcellular structures such as spines, and finer voltage changes. In densely packed populations of GEVI-expressing neurons, the high brightness of JEDI indicators at resting potential can introduce significant photon shot noise from neuropil background fluorescence. In this context, detecting spike responses, characterized by rapid fluorescence decreases, becomes more challenging. Reversing the response polarity of these indicators could enhance spike detection by improving contrast against the baseline, given larger spike responses amplitude and substantial baseline fluorescence levels^50^. However, such “positive-going” indicators may exhibit insufficient baseline fluorescence, reducing detection of hyperpolarizations and small depolarizations. These trade-offs highlight the need for a diverse collection of GEVIs optimized for specific experimental needs.

JEDI3 sensors significantly expand the two-photon imaging toolbox, enabling more precise detection of subthreshold voltage changes. Future GEVI enhancements, combined with advances in optical techniques and denoising algorithms, will increase the robustness and versatility of voltage recordings, broadening their applicability to new experimental paradigms. JEDI3 indicators have already facilitated experiments that were previously challenging or impossible, providing novel insights into neuronal activity at both subcellular and population levels. By integrating JEDI3 sensors with emerging imaging technologies and optogenetic approaches, researchers can further unravel the intricate relationships between subthreshold voltage dynamics, neurotransmitter signaling, and circuit-level processes. This work opens new avenues for discovery in neuroscience, facilitating the study of normal brain function and the pathophysiology of neurological disorders.

## Methods

### High-throughput GEVI screening

#### Single plasmid and library construction

Plasmids used as controls or for *in vitro* characterization were assembled using the In-Fusion cloning technique (Takara Bio USA, Inc.) into a pcDNA3.1/Puro-CAG vector with or without a C-terminus GSSGSSGSS linked mBeRFP E6F reference protein^51,52^. The plasmid sequence was confirmed using whole plasmid sequencing based on nanopore technology (Plasmidsaurus Inc.) or by Sanger sequencing (Eurofins Genomics LLC).

Libraries were constructed as previously described using degenerate codons and In-Fusion cloning to assemble the library^8^ with slight modifications. For library screening variants were GSSGSSGSS linked with mBeRFP E6F. For PCR reactions, we decreased the amount of primer and template used in the PCR reaction to 1 µl of 10 µM and 5 ng, respectively. The amplification cycles in the PCR reaction were reduced to 30. Liquid cultures were inoculated with manually picked colonies, and purified plasmids were prepared using a 96-well plasmid <u>purification</u> kit (96 wells Mini Plus Plasmid Extraction System, Viogene) following the manufacturer’s instructions.

#### In silico modeling

3D visualization of JEDI3 sensors used a modified 3D structure previously generated^8^. Visualization and labeling were done using PyMOL.

#### Cell culture and transfection in 96-well plates

HEK293 cells stably expressed human Kir2.1 channel^53^ were used to screen for sensors with a resting membrane potential at approximately −83 mV in our conditions. Cellular growth conditions and transfection protocols were as previously described^8^. We generally selected 88 variants per library. According to a statistical model, our library generation and sampling strategy produced a ∼99% theoretical probability that any given library included the best residue^54^. The imaging solution was the same, except that the final HEPES concentration was increased to 20 mM.

#### Two-photon screening system

Two-photon library screening was completed with the same microscope, laser, objective, filters, PMTs, stimulator and PC used for the engineering of JEDI-2P^8^. Images were acquired via resonance scanning at about 440Hz. The fluorescence signal was collected from both the GEVI and the mBeRFP E6F reference protein. Each FOV (512x32) was illuminated with 940nm light tuned to 75 mW at the sample plane. All stimulations were monophasic square pulses and the protocol started with 5x60V pulses for 1 ms with an inter-pulse duration of 300 ms. These 5 stimulations were then followed 1.3 sec later by a ramp that varies in voltage and time with the sequence 15 V for 1 ms, 25 V for 1 ms, 30 V for, 30 V for 2.5 ms, 30 V for 3.5 ms, 30 V for 4.5 ms, and 30 V for 7.5 ms. The onset of the ramp stimulations were separated by 300 ms. The ramp was followed by a 100 Hz train comprised of 10 pulses at 30V for 2.5 ms. Concluding the stimulation protocol was 5 more pulses at 60V for 1 ms with an inter-pulse duration of 300 ms.

Images were analyzed using the same custom routines in MATLAB (version R2024a, MathWorks) as was used for the identification of JEDI-2P^8^. Photobleaching was quantified as the area under the fluorescence trace after it had been normalized to fluorescence at t = 0. Peak response amplitude was quantified from the photobleached corrected fluorescent trace and was the average across 4 FOV from 1 well. If there was more than 1 well per variant on a plate each well was then be averaged together. GEVI brightness was calculated using the averaged fluorescence intensity of the first 20 frames in the green channel normalized by the average fluorescence intensity of the first 20 frames in the red channel. Using the same custom compound metrics developed previously we were able to rank sensors while considering multiple performance criteria.

The detectability index is defined as response amplitude times the square root of the relative brightness. The detectability budget is defined as the detectability index times the square root of the photostability.

### GEVI characterization *in vitro*

#### Preparation for voltage clamp

Cell culture and transfection protocols are the same as previously described^8^ with minor modifications. HEK293A cells (Thermo Fisher Scientific) were plated on a round coverslip (12 mm #0, 633009, Carolina) in a 24-well flat bottom culture plate (Costar, # 3524). On the same day, cells were plated and transfected using 100 ng DNA and 0.3 μL FuGENE® HD (Promega Corporation) transfection reagent per well of the 24-well plate following manufacturer’s instructions. Unless indicated otherwise, pcDNA3.1/Puro-CAG plasmids expressing the GEVI with no reference protein were used. Media was changed the next day and imaged two days post transfection. Glass micropipettes (1B150-F-4, World Precision Instruments) were pulled (P1000, Sutter) to achieve a tip resistance of 2-6 MΩ. The internal solution and the external solution were the same as previously described^8^ with the exception that the external solution now had 20 mM HEPES. The coverslip seeded with the transfected cells was placed in a custom glass-bottom chamber based on Chamlide EC (Live Cell Instrument) with a square glass bottom (24 × 24 mm #1 coverslip, Thermo Scientific). The hardware setup was the same as previously described for whole-cell patch clamping. Command voltage waveforms were compensated for the liquid junction potential of HEK293A cells. Cells were included in the final analysis only if the patched cell had an access resistance (Ra) less than or equal to 20 MΩ and a <u>membrane resistance</u> (Rm) larger than 10 times of Ra both before and after the recording.

#### Preparation for current clamp

The setup for the current clamp was the same as the voltage clamp with slight modifications. The resting membrane potential of HEK293-Kir2.1 cells was determined using a whole-cell current clamp in the same imaging solution used for the 96-well plate screening and compensated for the liquid junction potential. HEK293-Kir2.1 cells were not transfected and plated the day before patching.

#### Voltage clamp under two-photon illumination

Voltage clamp experiments were done with the same protocols, equipment and analysis methods as previously described with some modifications^8^. All two-photon electrophysiology was done at 22-23°C, with no reference protein and illuminated with 940nm light at about ∼61 mW at the sample plane. Cells were stimulated with 5 AP waveforms with a FWHM of 2 ms at 2 Hz, 5 AP waveforms with a FWHM of 4 ms at 2 Hz and 10 AP waveforms at 100 Hz. In addition, we also modified a burst of APs recorded from the adult mouse somatosensory cortex L5 pyramidal neurons to mimic APs on top of subthreshold depolarizations. The spike burst has a subthreshold depolarization of ∼24OmV (from –70OmV baseline voltage) and APs of 60–90OmV amplitude (from –56OmV subthreshold voltage). The APs in the spike burst are 3–4Oms FWHM. The last part of the protocol held the cells for 1s at different voltage steps from a holding potential of -70 mV. Those voltage steps were -120, −100, −80, −60, −40, −20, 0, 20, 30, and 50 mV, followed by more refined steps around subthreshold of -80, -85, -85, -60, -55 and -50 mV. After the AP waveforms a 2 sec interval at -70mV was applied before the start of the voltage steps and then a 1.5 sec interval between each voltage step. Videos were taken with a resolution of 512 × 32 pixels and a frame rate of 440 Hz.

#### Brightness at -70 mV using stably expressing cell lines

Using transient transfection techniques produces a population of cells that have large cell-to-cell variation due to differences in plasmid copy number and translational differences^55^. To compensate for plasmid copy variation, we developed a stable cell line using the Flip-In^TM^ HEK293 cells (Invitrogen, #R75007), which enable single copy number chromosomal integration. To compensate for differences in translation, we added a C-terminus GSSGSSGSS linked mBeRFP E6F reference protein. JEDI3hyp, JEDI3sub and JEDI-2P with mBeRFP E6F were cloned into a pcDNA5/FRT backbone via previously described cloning methods. Flip-In^TM^ HEK293 cells were plated 4 hr before transfection at 70% confluency into a 96-well plate (Corning, #353916). Cells were transfected with premixed 20 ng of the pcDNA5/FRT GEVI and 180ng of the pOG44 plasmid. DNA was mixed with 0.6 ul of room temperature FuGENE® HD. Then 13 ul of optiMem (Thermo Fisher Scientific, #11058021) was added to the FuGENE and DNA mixture. The reaction was allowed to incubate at room temperature for 8 min. After incubation, 50 ul of growth media (same as used for cell growth of HEK293A and HEK293 Kir2.1 cells) was added to the reaction. The total volume was transferred to each well. After 24 hr of transfection, cells were washed with DPBS (Corning, #21-031-CV), and fresh growth medium was added. After 48 hrs of transfection cells were split into 4 new wells of a 96-well plate at a confluency of around 25%. During this split, the hygromycin (Gibco™ Hygromycin B, #10687010) selection reagent was added at a concentration of 25 µg/ml. Cells were cultured and every 3-4 days selective medium was changed. After 20 days the cells were considered to be stably expressing the JEDI3hyp, JEDI3sub and JEDI-2P sensors. The cells were cultured following previously described culturing protocols with the addition of 25 µg/ml of maintenance Hygromycin.

Brightness was determined at -70 mV using electrophysiology protocols and equipment as described in this paper. Cells were held at -70mV for 30 seconds, and images from both the green and red channels were collected. Cells were illuminated with 940nm that was 74 mW at the sample plane. Images were taken using two galvanometer optical scanners at 1.8 frames per second with 2-frame averaging. Saturated pixels, typically from over-expressing cells, were excluded and background levels were estimated using the intensity histogram and subtracted from images for both channels. An initial mask was computed from the first frames of each channel using predefined thresholds to enrich for pixels associated with cell membranes. This mask was applied to the entire recording. The fluorescence intensity for the first 20 images were then averaged for both the red and green channels. The mean of the green channel was divided by the mean of the red channel to give the relative brightness at -70 mV for each sensor.

#### Voltage clamp under one-photon illumination

The same microscope was used as previously described^8^. Cells were illuminated with 470/24-nm light (SpectraX, Lumencor) and conditioned using the 477-503-nm band of a multi-band dichroic mirror (89100bs, Chroma). The irradiance at the sample plane was ∼4 mW/mm^2^. Green emitted photons were reflected towards the camera or PMT using the 503-542-nm band of the multi-pass dichroic (above) and filtered at 509-532 nm using a multi-pass filter (89101m, Chroma). Kinetics were evaluated at 32-33°C by using three 1 sec 100-mV depolarization pulses from −70 to 30 mV. To characterize kinetics at different voltage steps corresponding to hyperpolarization, subthreshold and APs we used 3x1 sec steps (total 9 steps) from -70 to -100 mV, -70 to -45 mV and -45 to +30 mV. Between each pulse, cells were held at –70 mV for 1.4 s. The perfusion chamber and inline heater were the same as previously described^8^. To ensure only the patched cell was imaged a diaphragm was used to reduce the diameter of the excitation to just one cell. The same PMT and software was used to collect and analyze the data as previously described^8^. The three steps for each type of simulation are averaged. The photobleaching corrected mean signal was cropped from 0.1 sec before the estimated depolarization or the repolarization onset to 1 sec after the estimated depolarization or repolarization onset, with the exception of the -45 to -30 mV repolarization step, which was cropped 0.2 sec after the estimated repolarization onset. The exact onset timing was fitted together with other coefficients with either single-exponential (F(t) = c + (k × exp((t - t0) × λ)) × (t > t0) + k × (t ≤ t0)) or dual-exponential (F(t) = c + (k × exp((t - t0) × λ) + k2 × exp((t - t0) × λ2)) × (t > t0) + (k + k2) × (t ≤ t0)) model where the t is the independent variable, F is the dependent variable, and the rest are the coefficients to be fitted. Among these coefficients, c describes the mean plateau fluorescence, k or k2 describe the relative ratio of each exponential component, λ or λ2 describe (minus) inverse of the time constant(s), and t0 is an offset indicating the exact event onset timing. In some instances, λ2 describes a slow photobleaching component. For the -100 to -70mV and the -70mV to - 100mV steps, all three JEDI sensor kinetics were best fit by a dual-exponential where λ describes the off-kinetics and λ2 describes a slow photobleaching component not reported in Table 1.

#### Two-photon excitation spectra

To determine the two-photon excitation spectrum for JEDI3 sensors and controls we cloned them into the pcDNA3.1/Puro-CAG plasmid between the NheI and HindIII sites with no reference protein attached. HEK293 Kir2.1 cells were transfected using the same protocol as the above 96 well plate transfections. Images were acquired using the same microscope as described for two-photon screening. Laser pulses were not pre-compensated for dispersion in the microscope optical path. Excitation wavelengths from 700 to 1080 nm were used in 10-nm increments and tuned to a power of 16-18 mW at the sample plane, as measured by a microscope slide power sensor (S170C Thorlabs). Each FOV was imaged at each wavelength using two galvanometer optical scanners set to acquire a frame in 1.1 sec. Fluorescence values were corrected by subtracting the background, and small deviations in the power between wavelengths were corrected by assuming a quadratic dependence of fluorescence on illumination power at the sample plane. The same power was applied for each sensor and control.

#### Two-photon photostability characterization under different powers

Using the same hardware and data processing methods as previously described^56^, we were able to characterize the JEDI3 variants with some modifications. Cells were cultured and transfected with JEDI3 sensors and JEDI-2P without reference protein in 96-well plates, as discussed in the Cell culture and transfection in 96-well plates section. Cells were imaged in the same external solution as previously described. Photobleaching was conducted under 940nm with power ranging from 29 to 101 mW at the sample plane. Each well had 6 separate FOV 512x32 pixels that were continuously imaged for 6 minutes at 440 Hz. The emission light from the cell was split using a 560-nm dichroic mirror (348958, Chroma), filtered by a 525/50-nm bandpass filter (353716, Chroma) and collected by a gallium arsenide phosphide (GaAsP) photomultipliers tube (PMT).

To calculate the area under the curve for each GEVI, the videos were first background-corrected and foreground-segmented using a predefined threshold as described in the brightness at -70 mV section. The averaged fluorescence from the foreground pixels was normalized to the fluorescence of the first frame of the video. The area under each curve was then calculated and averaged for each variant. A two-way ANOVA with Dunnett’s multiple comparisons test was used to determine significance. The multiple comparisons test was to compare the JEDI3 sensors to JEDI-2P for each power level. The fitting of the 37 mW fluorescence traces was fit by an exponential equation modeling a two-phase decay.

#### Confocal imaging of GEVI in dissociated neurons

E18 mouse hippocampal neurons were purchased from Transnetyx tissue by BrainBits (SKU SDEDHP, Transnetyx, Inc.) and came in Hibernate® EB complete Media as single-cell suspensions. Prior to seeding the hippocampal neurons, a 24-well glass bottom plate (P24-1.5H-N, Cellvis) was coated with Neuron Coating Solution (027-05, Sigma-Aldrich) at 37O°C overnight and washed three times with PBS. Neurons were seeded at 80,000–100,000 cells/mL in 500OµL glutamate-supplemented NbActiv1 medium (NB1O+OGLU, Transnetyx Inc.) per well of the 24-well plate following the manufacturer’s protocol. The plating day was considered as DIV 0. Every 3-4 days half of the media was replaced with pre-equilibrated NbActiv4 medium (SKU NB4, Transnetyx Inc.). Transfection was conducted on DIV 12–14 using 1OµL lipofectamine 2000 and 800Ong total DNA, a mix of 150Ong pAAV-hSyn-JEDI3hyp or pAAV-hSyn- JEDI3sub and 650Ong pNCS bacterial expression vector as buffer/filler DNA.

Two days after transfection (DIV 14–16), the attached neurons were washed twice with the same external solution used in the two-photon screening system before adding a final 500OµL per well as the imaging solution. Laser-scanning confocal images were obtained using a high-speed confocal microscope (LSM880 with Airyscan, Zeiss) driven by the Zen software (version 2.3 SP1 FP3 black edition, Zeiss). The microscope was equipped with a 40× 1.1 NA water immersion objective (LD C-Apochromat Korr M27, Zeiss), a 488-nm argon laser (LGK7812, Lasos) set to 5% power (10OµW) and a per-pixel dwell time of 1.51 μs (JEDI3sub) or 3.36 μs (JEDI3hyp). Emission light was filtered using a multipass beamsplitter (MBS 488/561/633, Zeiss) and acquired with a 32-channel GaAsP detector (Airyscan, Zeiss) with a detector gain of 747 (JEDI3sub) or 800 (JEDI3hyp), and a 1.63 (JEDI3sub) or 1.08 (JEDI3hyp)-Airy unit pinhole size. Images were acquired at a resolution of 0.04Oµm/pixel and an X-Y dimension of 2752 × 2752 pixels (JEDI3sub) or 4972 × 4972 pixels (JEDI3hyp). Z stacks were stepped at 7Oµm (JEDI3sub) or 4 µm (JEDI3hyp) between images. To increase the signal-to-noise ratio, 8 scans for JEDI3sub and 4 scans for JEDI3hyp were performed and averaged for each image. Airyscan processing was applied to the images via Zen software (version 2.3, blue edition, Zeiss) to increase the resolution. Fig. 1B corresponds to a maximum intensity projection from Z-stack with 8 images for JEDI3sub or 5 images for JEDI3hyp. The overall image and dendrite zoom-in are from the maximal projection of the z stack, whereas the soma image is a single slice.

### Cloning and packaging AAVs for *in vitro* experiments

All cloning was performed in-house using In-Fusion cloning techniques discussed in the high-throughput screening methods section. Using the pAAV-EF1a-DIO-JEDI-2P-Kv-WPRE plasmid (Addgene, #179459) we cloned JEDI3hyp and JEDI3sub in place of JEDI-2P. The Kv tag promotes enrichment of the GEVI to the soma^8,27^. Unless otherwise stated, the double-floxed inversed JEDI sensors under the control of EF-1α promoter was then packaged into Adeno-Associated Viruses serotype 1 (AAV2/1) at the BCM Neuroconnectivity Core or the CNP Viral Vector Core at the CERVO Research Center contribution (RRID:SCR_016477).

### Optical recordings in mouse cortex using ULoVE

All protocols adhered to the guidelines of the French National Ethics Committee for Sciences and Health report on Ethical Principles for Animal Experimentation in agreement with the European Community Directive 86/609/EEC under agreement #29791.

### Animal handling, viral injections, and surgeries

8 male wild-type C57BL/6J adult mice (>P40 - body weight 20–24 g, n=3 for JEDI-2P, n=2 for JEDI3hyp and n=3 for JEDI3sub) were housed in standard conditions (12-hour light/dark cycles, light on at 7 a.m., with water and food ad libitum). A preoperative analgesic was used (buprenorphine, 0.1 mg/kg), and Ketamine-Xylazine was used as an anesthetic (Centravet). The AAV2/1-eF1-DIO-JEDI-2P-Kv-WPRE virus was made in-house^57^ using a plasmid ratio (µg) of transgene:capsid:helper: 1:1.6:2. The AAV2/1-EF1a-DIO-JEDI-2P-Kv-WPRE, AAV2/1-EF1-DIO-JEDI3hyp-Kv-WPRE and AAV2/1-EF1-DIO-JEDI3sub-Kv-WPRE viruses were injected at a titer of 3.10^12^ GC/ml. To obtain a sparse density of GEVI-expressing neurons, AAV2/1 hSyn-Cre (Addgene, AV-1-PV2676) was co-injected at a final titer of 2.10^9^ GC/ml).

Viruses were combined in a saline solution containing 0.001% of pluronic acid (ThermoFischer 24040032), 300 nl of which was injected at a flow rate of 75 nl/min into the visual cortex (V1 coordinates from bregma: anteroposterior –3/–3.5 mm, mediolateral –2.5/–3 mm, and dorsoventral –0.3 mm from brain surface). A custom-designed aluminum head-plate was fixed on the skull with layers of dental cement (Metabond). A 5-mm diameter #1 coverslip was placed on top of the visual cortex and secured with dental cement (Tetric Evoflow). Mice were allowed to recover for at least 15 days before recording sessions and housed at least 2 mice per cage. Imaging was done between 15 and 77 days post-surgery. Behavioral habituation was adopted, involving progressive handling by the experimenter with gradual increases in head fixation duration^58^. Mice were handled before recording sessions to limit restraint-associated stress, and experiments were performed during the light cycle.

### UloVE voltage optical recording and experimental design

Recording sessions were 1-3 hours long and were performed while mice behaved spontaneously on top of an unconstrained running wheel in the dark^58^. Recordings were performed using a custom designed acousto-optic deflectors (AOD) -based random-access multiphoton system (Karthala System) based on a previously described design^17^. The excitation was provided by a femtosecond laser (InSight X3, Spectra Physics) mode-locked at 920 nm with a repetition rate of 80 MHz. A water-immersion objective (CFO Apo25XC W1300, 1.1 NA, 2 mm working distance, Nikon) was used for excitation and epifluorescence light collection. This experiment was done at 920nm because the peak 2P excitation wavelength for the JEDI3 sensors had not yet been determined. The signal was passed through an IR blocking filter (TF1, Thorlabs), split into two channels using a 562 nm dichroic mirror (Semrock), and passed to two H12056P-40 photomultiplier tubes (Hamamatsu) used in photon counting mode. The 510/84 filtered green channel was used for collecting JEDI signals. For imaging preceding ULoVE recordings, the same AOD based microscope was used but in imaging mode. A time series of 50 images (1µs/pixel, 0.091µm/pixel) was acquired and post hoc motion registered to produce the images in Fig. 2B. Cell depth was measured from the surface of the brain.

The optimized ULoVE excitation pattern (Lombardini et al., in prep) consists of a series of 9 points vertically aligned (evenly spread over a distance of 15µm), multiplexed twice horizontally with a 2µm spacing. The 18 points are scanned in diagonal (8 µm horizontally and 3µm vertically) during the acquisition time of 50 µs, in order to homogeneously fill an extended excitation volume continually encompassing the cell plasma-membrane. This latest strategy refines the axial profile of excitation, limiting signals out of the desired focal plane, thus reducing any neuropil signal contamination and improving SNR. For imaging, laser power was set to deliver 15 mW post-objective and pre-sample then adjusted for mono-exponential loss through tissue with a length constant of 170 µm. For ULoVE optical recordings, we further multiplied the power by 1.5, compared with the applied power used in the imaging mode, to account for the greater excitation volume. The applied power never exceeded 200 mW. Using two patterns per cell enabled a temporal resolution of 7142.9 Hz. Recordings were stopped after 320 seconds (5.33 minutes). For recording, we selected neurons that were sufficiently bright to obtain significant signal-to-noise, yet did not display long-lasting depolarizing plateaus, indicative (in our observations) of over-expression.

### Signal analyses, spikes extraction and waveform analyses

For each cell recording, two ULoVE volumes were simultaneously used as “regions/volumes” of interest (Fig. 2B), thus traces represent the addition of photons collected from the volumes. Photobleaching was assessed by bi-exponential fitting and corrected by division of the raw trace by the normalized fit function. This analysis enabled us to account for the fast-photobleaching component and correct the full trace used for further analyses (see below). To characterize the second phase (from a minute till the end), which could be more the consequence of a diffusion process described by a power law, we low-pass filter the trace at 1Hz, then down sample this trace 140 times and performed a linear fit on log10 log10 scales allowing us to get the slope of the power law.

To generate the %DF/F trace, we took the raw trace normalized by the bi-exponential fit function (computed above), after removing the remaining low frequency drift using a zero phase distortion filter (high pass: 0.5 Hz), the trace was converted to % ΔF/F taking the mean signal as F_0_.

Spikes were detected using a custom designed algorithm, which utilizes three metrics to sort aspects of spike shape and threshold, all with Zscores above 3. The first metric is to high pass filter the trace (second order Butterworth filter with lower limit set at 40Hz). The second metric is a cumulative probability transformation of the signal using the standard erf function for a duration of 1.6 ms. The third metric takes the cumulative product of the 2^nd^ to the 5^th^ scale of the coif1 discrete wavelet transform and applies a global realignment to retrieve energy in various frequency bands in one peak.

To extract the spike waveform metrics for quantifications (Fig. 2F), an average spike waveform was aligned on the spike onset. Then spike amplitude was taken as the peak value of the spike waveform average, measured from the onset point. FWHM corresponds to the extent in time at half maximum amplitude, taken from the average spike waveform after 20 kHz linear interpolation.

To quantify subthreshold fluctuations, traces were filtered using a bidirectional butterworth bandpass filter between 0.1 and 50Hz. The cell’s down-state corresponds to the 1^st^ percentile of that filtered trace. The cell’s up-state corresponds to the average of the signals at the detected spike onsets. The subthreshold fluctuation is taken as the difference between the cell’s up-state and the cell’s down-state. The dynamic range is the sum of the spike amplitude and the subthreshold fluctuation.

### JEDI3sub tracking of population-level subthreshold optical tuning

All animal experiments were conducted according to the National Institutes of Health guidelines for animal research. Procedures and protocols on mice were approved by the Institutional Animal Care and Use Committee at University of California, Berkeley.

### Craniotomy and virus injection

Virus injection and cranial window implantation procedures have been described previously^30^. Briefly, two mice (94 days old) were anesthetized with isoflurane (1-2% by volume in oxygen) and given the analgesic buprenorphine (subcutaneously, 0.3mg per kg of body weight). Animals were head fixed in a stereotaxic apparatus (Model 1900, David Kopf Instruments). A 3.5-mm-diameter craniotomy was made over the left visual cortex with dura intact. Wild type mice (C57BL/6J) were used, and virus injection was performed to express pAAV-hSyn-JEDI3sub-Kv2.1 (1.17×10^12^ GC/mL, 10× dilution), using a glass pipette with a 15-20 μm opening and a 45° bevel. The pipette was backfilled with mineral oil, and a fitted plunger controlled by a hydraulic manipulator (Narishige, MO10) was inserted into the pipette for loading. Virus was injected slowly at 6-12 sites (50 nl at each site) at a depth of 250 μm within the left visual cortex. A glass window made of a single coverslip (Fisher Scientific No. 1.5) was embedded in the craniotomy and sealed in place using dental acrylic. Then a titanium head-post was affixed to the skull using cyanoacrylate glue and dental acrylic. *In vivo* imaging was conducted after 3 weeks of recovery. All imaging experiments were carried out on head-fixed awake mice.

### *In vivo* population voltage imaging

We imaged the population subthreshold voltage activity from L2/3 neurons with a subkHz frame rate two-photon fluorescence microscope (2PFM)^25,29^. Output from a 1035 nm fiber laser (Monaco 1035-40-40, 1 MHz repetition rate; Coherent) was focused by a cylindrical lens and then sent into a pair of mirrors with a small tilt angle. Based on free-space angular-chirp-enhanced delay (FACED)^59^, the laser pulse was split into multiple pulses that were spatially separated and temporarily delayed by the mirror pair, forming a 1D array of excitation foci at the objective focal plane with a line-scanning rate of 1 MHz. A galvanometer scanned excitation foci in directions perpendicular to the linescans to form 2D images, and an additional galvanometer tiled the 2D images to cover a larger FOV. With a 25× 1.05 NA objective (XLPLN25XWMP2; Olympus), we imaged L2/3 neurons in the mouse visual cortex at depths ranging from 110 µm to 160 µm over 400 µm × 320 µm or 400 µm × 160 µm FOV at a frame rate of 400 Hz or 769 Hz, respectively, at a post-objective excitation power of 156 mW.

### Visual stimulation

A custom script written with the Psychophysics Toolbox^60^ in MATLAB (Mathworks) was used to display on an LCD monitor drifting grating visual stimulation to the right eye of the mouse. Each visual stimulation trial consisted of 8 gratings drifting in angles from 0° to 315° in 45° intervals, with a 0.5-second blank period preceding each 0.5-second drifting grating. 24-26 trials were acquired for each imaging FOV.

### Population voltage image analysis

Images were analyzed using custom programs in MATLAB (Mathworks). Time-lapse images were motion registered using an iterative cross-correlation method^30^, with the time-averaged image from the first trial as the registration target for the remaining trials. ROIs were first selected using the automatic cell segmentation pipeline Cellpose 2.0^61^ and were then manually curated to add new ROIs and remove incorrectly identified regions. Then, the average signal within each ROI was extracted as the raw neuronal trace F_raw_. To calculate ΔF/F, the baseline signal (F) was obtained by averaging the signal from 0.3 to 0.5 s during the blank period preceding each grating stimulus, and ΔF was defined as F_raw_ – F, the difference between the raw trace F_raw_ and the baseline F. To calculate the subthreshold activity, the raw -ΔF/F trace was low-pass filtered with a 3rd-order, 12-Hz Butterworth filter. The mean -ΔF/F value during the 0.5-s stimulation period and the 0.3-s blank period following the stimulation period was defined as the subthreshold response to the stimulation.

Even though the Kv tag reduces JEDI3sub expression in the neuropil, we carried out neuropil subtraction^62,63^. Following a common convention for neuropil subtraction in calcium imaging^64^, the fluorescence trace for each ROI was calculated as F_ROI_ = F_raw_ - 0.7*F_neuropil_, with F_neuropil_ being the average fluorescence of pixels within a neuropil mask surrounding each ROI. To get the neuropil mask, a ring was grown around the ROI, and the growth stopped when the mask size became more than 300 pixels. Pixels from other ROIs, and pixels with extremely high intensity values (top 4% of the whole field-of-view, corresponding to puncta) were removed before generating neuropil masks. On average, the neuropil mask size was 1.4 times the size of the ROI. ROIs with F_neuropil_ >1.1*F_raw_ were excluded before further analysis.

To determine whether a cell had visually evoked (VE) subthreshold activity, a tailed t-test was performed for each drifting direction to compare whether the subthreshold response to the stimulation (as defined above) was significantly larger than the average -ΔF/F during the final 0.2 s of the preceding blank period. If the p-value was less than 0.0063 (0.05/8, correction for multiple comparison), then it was considered a VE cell. A VE cell was considered to have orientation selectivity (OS) if the p-value from a one-way ANOVA test, which assessed how subthreshold responses vary with the direction of drifting grating, was less than 0.05.

To quantify the orientation tuning properties of VE cells, its subthreshold response *R* to each stimulation orientation and stimulation orientation angle (*θ*) were fitted using a bimodal Gaussian function65:

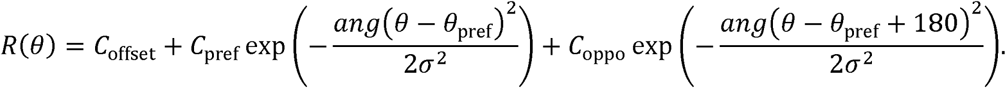

Here *θ*_pref_ is the preferred orientation, *θ*_pref_ and *θ*_oppo_ are the response coefficient at *θ*_pref_ and *θ*_pref_ 180, respectively. The *ang*(*θ*) wraps *θ* to the interval of 0° to 180°. The *σ* is the standard deviation of the Gaussian function. The global orientation-selectivity index (gOSI) of the subthreshold response is defined as ^30^:

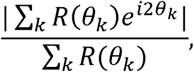

where *k* is the orientation index

Correlations between cell pairs were analyzed by calculating the Pearson’s correlation coefficient of the responses between each cell pair^66^.

### JEDI3hyp tracks sharp-wave ripples in PV hippocampal interneurons

All procedures were carried out in accordance with NIH guidelines and with the approval of the IACUC of Baylor College of Medicine.

### Animal handling, viral injections, and surgeries

Four adult mice (8 weeks or older at start of experiment) were used. Mice were housed in groups of 2-5 animals, on a normal diet and 12/12 light/dark cycle. PV-Cre mice were obtained from JAX (#017320) and were bred as homozygous pairs.

Mice were anesthetized with isoflurane and treated with analgesics (bupivacaine and buprenorphine). The skull was exposed, and a hole was drilled above the right dorsal hippocampus. Mice were injected in the dorsal CA1 (2.3mm posterior, 1.5 mm lateral, 1.45-1.35mm ventral to Bregma) using a 1 microliter Neuros (Hamilton) syringe, with AAV2/1-EF1-DIO-JEDI3hyp-Kv-WPRE (400 nl each, full titer at 4×10^12^ GC/mL). Injection was started ventrally, and virus was delivered in pulses of 50-100 nl while gradually raising the syringe within the dorsoventral range, with 1-minute intervals between pulses.

In order to implant chronic imaging windows and headbars, surgeries were performed after five days or more of postoperative recovery following viral injections. Mice were surgically implanted with an imaging window made of a round glass coverslip (diameter:3 mm, Warner), attached to a stainless steel cylinder (diameter: 3.18 mm, McMaster-Carr) with Narland optical adhesive. Mice were anesthetized with isoflurane and treated with analgesics (bupivacaine and buprenorphine), the skull was exposed, and a 3.0 mm diameter circular craniotomy centered on the injection site was marked with a robot stereotax (Neurostar) and finished with a hand-held drill (Osada). Dura and cortical layers were removed by vacuum suction while flushing the brain with ice-cold cortex buffer^67^, to minimize bleeding. The cannula was inserted and secured with tissue adhesive (Vetbond). The cannula and the exposed skull were covered with self-etching UV-curable adhesive (Optibond), and a stainless steel headbar was attached with dental cement (Teets). After postoperative recovery, mice were returned to home cages in the vivarium.

After recovery from surgery and confirming clean optical access to the hippocampus (10-14 days after surgery), mice were implanted with chronic hippocampal electrodes. A craniotomy was made, and a twisted tungsten wire pair made with 0.5 mm tip separation (0.002”, A-M systems) was lowered in the dorsal CA1 mirroring the virus injection site. The probe was stabilized using UV-curable glue, and the connector pins (A-M systems) were attached to the headbar implant with dental cement.

### *In vivo* two-photon voltage imaging and experimental design

*In vivo* imaging at 940nm was performed on a setup consisting of the following: a two-photon, 8 kHz resonant scanner microscope (Bruker Investigator); a pulsed infrared laser (Spectra Physics MaiTai eHP-DS); a 16x objective (0.8 NA WI, Nikon). Laser power was controlled by a Pockels cell and was set to 100-200 mW (measured at the objective) depending on sample brightness. Image acquisition was controlled by Prairie View (Bruker). Image size was configured to 48 lines per frame, 256 pixels per line, resulting in 6 laser pulse/pixel sampling, and 288.8 Hz frame rate. Pixel size was configured anisotropically (0.94 by 1.87 microns in the x and y dimensions, respectively), to increase field of view size relative to the frame rate. Recording sessions from the CA1 oriens or pyramidal layers lasted 100,000 frames (5.8 minutes), during which the mice were freely running or resting on a 2 m-long, 2” wide cloth belt supported on bearing rollers on a custom treadmill. Treadmill position was recorded by rotary encoder connected to a PyControl system synchronized to the image acquisition system^68^. Local field potentials were recorded as a differential between both wire tips using a differential amplifier (Warner DP-304, 0.1 Hz to 1 kHz, 1000x gain) connected to the digitizer of the microscope system, and digitized at 2 kHz.

### Image and LFP signal processing

Movies were pre-processed, cells were segmented, and ΔF/F traces were computed in Python. Rigid motion correction was performed using suite2p^69^ then the following steps were performed using a custom Python pipeline (https://github.com/bdudok/LNCM-lib). 1) polygon regions of interest (ROIs) were drawn manually on the average images; 2) mean pixel intensity was computed in each frame in each ROI; 3) the mean pixel intensity trace from a manually drawn, neuron soma-sized background region (without noticeable labeled profiles) was subtracted from each cell’s pixel intensity trace, resulting in the background-subtracted time series (F); 4) a 3^rd^-degree polynomial was fit on F^36^, excluding frames when the mouse was running, and any fluorescence peaks over 2 standard deviations; to obtain the time-dependent baseline (F_0_); 5) -ΔF/F was computed by subtracting (F-F_0_)/F_0_ from its maximum. Smoothed traces (only used for visualization, not for quantification) were computed using an exponentially weighted moving average (ewm method of pandas).

Sharp wave-ripples were detected automatically from band pass-filtered LFP traces (130-200 Hz). The algorithm described earlier^34^ was implemented in Python using the bessel, filtfilt and hilbert functions of scipy.signals. Events were curated manually to exclude false positives, by an expert investigator blinded to imaging data. Event onsets were defined as the time of the Hilbert envelope crossing a threshold set at 3 standard deviations above mean. Ripple-band power in each imaging frame was calculated as the mean of the envelope during each sample measured during the frame, and the resulting time series was divided by the standard deviation.

SPW-R onset times were clustered using DBSCAN (eps = 1 s), and single events and the first events of clusters were included to calculate peri-event averages. Masks around the event onsets were determined such that each imaging frame was only included once in the average, even if consecutive events were closer than the investigated time window. Frames were only included when the mouse stayed immobile. Generation of average traces (fluorescence or ripple-band power) were calculated first from the cell-wise means, then the mean ± standard error (SEM) of the cell-wise means was computed using the number of animals as the sample size for calculating SEM. Response magnitudes were quantified by using the locations of the maximum and minimum of the post-ripple grand average (21 and 114 ms, respectively), then each individual cell’s intensity was averaged during a 50 ms time windows centered on the locations, finally the baseline of the cell (-200 to -100 ms window before ripple onset) was subtracted from the responses. To determine whether the responses were different from the baseline, statistical tests were done on the animal-wise averages, using scipy (1-sample, 1 sided t test, with alternative hypothesis “greater” for peak and “less” for trough).

### JEDI3hyp Two photon Z-stacks

*In vivo* two-photon Z-stacks (Fig. S4) were acquired from awake, head-fixed mice before any functional imaging, using the galvo-galvo modality of the two-photon microscope, with 1024x1024 resolution, 1.36 x 1.36 µm lateral pixel size and 5 µm axial step size. Images were converted to RGB format using ImageJ, and identically sized regions were cropped with Adobe Photoshop for each panel. Image contrast was enhanced using the Curves function of Photoshop, applied uniformly to all images.

### JEDI3hyp optical tracking of brain state in mouse cortex

All procedures were carried out in accordance with the ethical guidelines of the National Institutes and were approved by the Institutional Animal Care and Use Committee (IACUC) of Baylor College of Medicine. The following experiments in this section were done at 920nm because the peak 2P excitation wavelength for the JEDI3 sensors had not yet been determined.

### Animal handling, viral injections, and surgeries for *in vivo* 2P Imaging in Mice

We use data from a total of 13 mice aged 6 weeks to 6 months.

Anesthesia induction was performed with 3% isoflurane and maintained with 1.5% isoflurane throughout the surgical procedure. Mice were injected with 5 mg/kg meloxicam or ketoprofen subcutaneously prior to the start of the start of the surgery. Anesthetized mice were placed in a stereotaxic head holder (Kopf Instruments) with body temperature maintained at 37° C using a homeothermic blanket system (Somnosuite with RightTemp Module, Kent Scientific). The scalp is shaved and cleaned with alcohol pads and betadine after which 5 mL of 0.5% bupivacaine is injected subcutaneously under the scalp. After waiting 10 minutes, approximately 1 cm^2^ of skin is removed above the skull and along with underlying fascia. The edges of the incision are sealed with a thin layer of surgical glue (VetBond, 3M) followed shortly by the implantation of a 4 mm washer (Seastrom) to head fix the mouse using a custom, removable headbar. The washer was implanted using dental cement (C&B Metabond, Parkell) over our region of interest, visual cortex (2.5-3 mm lateral and 1 mm anterior of the intersection of the midline and lambda suture). The mouse was subsequently transferred onto a small platform and held stationary via the headbar. Using a surgical drill and 0.6 mm carbide burr, a ∼ 4 mm craniotomy was made in the center of the washer and subsequently the cortical surface was washed and kept moist with ACSF (125mM NaCl, 5mM KCl, 10mM Glucose, 10mM HEPES, 2mM CaCl2, 2mM MgSO4).

In VIP, SOM, and L2/3 experiments JEDI3 expression was restricted perisomatically using AAV2/1-EF1-DIO-JEDI3hyp-Kv-WPRE (tither: 1.4x10^12^ gc/uL). 500-1000 nL was injected using a nano-injection pump (WPI) approximately 200-400 µm depth in VIP-Cre (RRID: IMSR_JAX:010908), SOM-Cre (RRID:IMSR_JAX:013044) or non-Cre expressing mice (wildtype, WT) co-injected AAV1-CamKII-0.4-Cre-SV40 (Addgene viral prep # 105558-AAV). For Layer 5 somatodendritic imaging (Fig. 5E,F), a dual-recombinase approach was used in Tlx-Cre mice to ultra-sparsely label single L5 IT somata and their apical dendritic arbors. 50-250 nL of AAV1-EF1a-DIO-FLPo-WPRE-hGHpA (Addgene viral prep # 87306) diluted 1:20,000 to 1:100,000 were co-injected with AAV1-EF1a-fDIO-JEDI3hyp approximately 500-650 µm in Layer 5 intratelencephalic (IT), Tlx-Cre (RRID: MMRRC_041158-UCD), mice. Following injection of viral construct(s), the dura was removed, avoiding damage to the cortex, and the craniotomy was secured and sealed with a 4 mm coverslip (Warner Instruments) and surgical glue (VetBond). Following the surgery, mice were singly housed in the mouse colony under standard conditions (12 h light/dark cycles) with water and food *ad libitum*.

### 2P Voltage Imaging of L1, L2/3, and L5 Pyramidal Neuron Somata via Resonant Scanning

The same 32 *in vivo* neuronal recordings of intratelencephalic (IT) Layer 5 pyramidal neurons were obtained using two-photon resonant imaging across four Layer 5 (IT) mice (Fig. 5B-D and Fig. S5A,C). Imaging power of L5 somata was kept between 77-115 mW at the objective for depth ranges of 443-567 µm. Cells were imaged at frame rate of 20-462Hz. For Fig. S5B, we used six neuronal recordings of Layer 2/3 pyramidal neurons from two non-Cre expressing mice (wildtype, WT). We restricted our neuronal recordings to Layer 2/3 pyramidal neurons restricting the imaging focal plane to layer 2/3 depths (100-350 µm depth) and imaging power was kept between 15 and 25 mW. Two-photon resonant imaging was performed on a Thorlabs Bergamo resonant scanning microscope with excitation via a Ti: sapphire femtosecond laser (Chameleon Vision II, Coherent) operated at 920 nm. A 1.1 NA 25x objective lens was used for imaging (CFI75 Apochromat 25XC W, Nikon Instruments). Emission light was split by a dichroic mirror into two channels: the green channel used a 525/50 nm filter, and the red channel used a 625/90 nm filter, before being collected by two photomultiplier tubes. ScanImage (Vidrio) was used to control the microscope and acquire imaging data. Pupil and running activity were obtained concurrently with neuronal recordings across all mice and microscopes.

JEDI photobleaching traces (Fig. S5E) were acquired using the same resonant scanning hardware described above. JEDI3 traces were calculated from manually segmented masks of soma across Layer 1, Layer 2/3, and Layer 5 that were at least 15 minutes long. Power measurements for each trace were obtained after imaging and measured post-objective using a Thorlab digital power meter (PM100D) with a thermal power sensor (S175C) using the imaging parameters for each scan. Fluorescence traces were trimmed down to 15 minutes, interpolated to have the same number of samples, and filtered using a low-pass Butterworth filter with a 0.1 Hz cutoff. Each trace was normalized by dividing the mean fluorescence within the first 5 seconds of imaging. Fluorescence traces were then grouped depending on their depth or cell type, prior to computing the average and the standard error of mean for each group.

### Simultaneous 2P Voltage Imaging of the Layer-5 Soma and Layer-1 Dendrites of Pyramidal Neurons via Resonant Scanning on a 2P-RAM Mesocope

Data for Fig. 5F and Fig. S5D is from 16 *in vivo* neuronal simultaneous recordings of intratelencephalic (IT) Layer 5 pyramidal neurons and 44 of their matching dendrites across four Layer-5 (IT) mice. Imaging was performed using two-layer 2P resonant scanning using a Thorlabs 2P-RAM mesoscope with excitation light delivered via a Ti: sapphire laser (Tiberius, Thorlabs) tuned to 920 nm and a custom 0.6-NA objective. The overall magnification was around 10x. Imaging ROIs (20-40 µm^2^) were placed over the soma and one or more matching apical dendrites. Imaging of multiple planes were performed at various volume rates ranging from 20-70 Hz. Excitation power when imaging L5 somata was kept between 77-115 mW at the objective at depths of 443-567 µm, and 20-50 mW when imaging their matching dendrites at depths ranging from 10-150 µm. Following functional imaging of individual soma and dendrites high-resolution (1 pixel/µm) z-stacks with 80-120 steps were obtained with 5 µm spacing across the entire somatodendritic axis spanning 400-600 µm. The structural z-stacks were then imported into NeuromanticV1.6.3^70^ for digital reconstruction of individual neurons and their apical dendrites. Pupil and running activity were obtained concurrently with neuronal recordings across all mice and microscopes.

### 2P Voltage Recordings of SOM and VIP Interneurons using Random-access Microscopy

We acquired 111 *in vivo* neuronal recordings of VIP interneurons from five VIP-Cre mice and 83 neuronal recordings from 2 SOM-Cre mice. The imaging frame rate ranged from 396 to 1040 Hz. Only recordings that showed a significant difference (p<0.01) in JEDI3hyp fluorescence during pupil dilation relative to constriction determined by a Kruskal-Wallis test were analyzed for Figure I (n=42/111) and Fig. S5F (n=7/83). SOM-Cre and VIP-Cre mice were crossed on a C57Bl/6 background and their neuronal recordings were obtained using acoustic optical deflector (AOD) imaging. 2P random-access imaging was performed on a Femto3D Atlas Plug and Play AOD 2P microscope (Femtonics) with excitation via a Spark laser operated at 920 nm. A 0.8 NA 16X and 1.1 NA 25X objective lenses were used for AOD two-photon voltage imaging. Single and multiple VIP and SOM somas were simultaneously imaged using chessboard and ribbon 3D scan modes. Depending on the depth, the imaging power was kept between 20-70 mW. Pupil and running activity were obtained concurrently with neuronal recordings across all mice and microscopes.

### Behavioral Data Acquisition and Preprocessing

Following two-weeks of recovery, mice were placed on a freely moving treadmill and head-fixed using a custom headbar for two-photon imaging. An optical encoder (McMaster-Carr) was used to track treadmill rotation. Running periods were determined with a velocity threshold of 1 cm/sec for at least 1 second. Running periods less than 3 seconds were consolidated into a single running period.

A camera (Dalsa) with a 1’’ hot mirror (Thorlabs) was used to record pupil activity at 20 fps. Since the experiments were performed in the dark without visual stimulation, the mouse was illuminated with a monitor backlight or UV light throughout *in vivo* 2P imaging experiments. Subsequently, a DeepLabCut model identified and tracked pupil edges from which the pupil diameter was computed. We transformed these measurements to physical units using a pixel to mm conversion based on the typical size of the mouse eye as previously detailed in^44^. Pupil periods were defined as pupil periods greater than one second with mean dilation or constricting speed greater than 0.02 mm/second. Eyelid blinks or frames with poor contrast where the pupil is not tracked by DeepLabCut were considered NaNs. Pupil periods composed of more than 15% NaNs were not analyzed. Given our interest in subtle brain state changes occurring during quiet wakefulness, we excluded pupil dilation and constriction periods that occurring 3 seconds before, after, and during running periods. Custom MATLAB and LabView (National Instruments) code were used to control the acquisition and synchronization of imaging and behavioral data.

### Preprocessing of voltage imaging data

Functional voltage imaging data was motion corrected in the horizontal plane for x-, y- shifts with a two-step algorithm. Large scale motion was first corrected by taking the average template of the entire scan and computing the phase correlated between the average template and individual frames. Following motion correction of large horizontal shifts, smaller scale motion was corrected by averaging frames of various windows sizes (ranging from frames within 3 secs to 2000 frames, depending on which had the best performance) and computing the phase correlation between this local template to individual frames. Somatodendritic imaging scans with greater than 3 µm of x-/y- shifts in the somatic imaging plane or greater than 1 µm of x-/y- shifts in the dendritic imaging plane were excluded for subsequent analysis. Following motion correction, VIP, SOM, L5 IT soma and dendrites were manually segmented.

### Pupil Phase Binning: Fluorescence response amplitude, ΔF/F_0_

For pupil phase binning, JEDI3hyp fluorescence traces were computed from all pixels in the manual segmented masks, inverted, low-pass filtered using a zero-phase 5 Hz Hamming lowpass filter, and subsequently normalized to ΔF/F_0_ by subtracting and dividing each raw fluorescence trace by F_0_, where F_0_ is the rolling 5^th^ percentile of a 20-minute window. We then iterated through all non-running pupil periods and binned the ΔF/F_0_ during that pupil period by the phase of the pupil diameter (65 phase bins from – π to π) to align all pupil periods to a single canonical cycle of dilation and constriction. All pupil phase binned fluorescence values were averaged, and error computed using standard error of mean (SEM). All fluorescence pupil phase plots are mean ΔF/F_0_ ± SEM for each bin. Finally, this phase-binned signal was minimally smoothed with a rolling average of window size 3 and centered around 0 by subtracting the mean.

### Pupil Phase Binning: Δ 2-10 Hz Hilbert transform amplitude

To compute the pupil phase-related change in low frequency oscillations, JEDI3hyp traces were computed from all pixels in the manual segmented masks, inverted, and bandpass filtered using a zero-phase 2-10 Hz Hamming bandpass filter. The Hilbert transform was applied to these traces and the amplitude of the transform was filtered between 0.1-1 Hz to match the timescales on which pupil periods occur. The Hilbert transformation detects the change in the signal envelope from the 2-10 Hz filtered trace. This transformation does not require averaging over many recordings, simplifying the analysis and improving the detection of frequency change when compared with wavelets or Fourier transformation. We then iterated through all non-running pupil periods and binned the amplitude measurement by the phase of the pupil diameter (65 phase bins from – π to π) to align all pupil periods to a single canonical cycle. These phase-binned fluorescence values were averaged, and error bars were computed using standard error of the mean (SEM). All fluorescence pupil phase plots are mean ΔF/F_0_ ± SEM for each bin. Finally, this phase-binned signal was minimally smoothed with a rolling average of window size 3 and centered around 0 by subtracting the mean.

## Data availability

The GenBank accession numbers are https://www.ncbi.nlm.nih.gov/nuccore/PX122673 for JEDI3sub and https://www.ncbi.nlm.nih.gov/nuccore/PX122674 for JEDI3hyp. Plasmids used in this work or that are useful for other applications are available on Addgene (ID numbers 246616–246641). Raw data for main and Supplementary Figures have been uploaded to Zenodo (https://doi.org/10.5281/zenodo.17537897). Data that is too large to upload will be made available upon request.

## Acknowledgments

**St-Pierre lab.** We thank M. E. Dickson, J.M. Kirk, C.-W. Hsu and the Imaging & Vital Microscopy Core (Baylor College of Medicine, BCM) for training on the confocal microscope. The project described was supported in part by the Neuroconnectivity Core at Baylor College of Medicine, which is supported by IDDRC Grant Number P50103555 from the Eunice Kennedy Shriver National Institute of Child Health & Human Development. The content is solely the responsibility of the authors and does not necessarily represent the official views of the Eunice Kennedy Shriver National Institute of Child Health & Human Development or the National Institutes of Health. **Bourdieu lab**. We thank Stéphane Dieudonné for invaluable advice on data production, analysis and interpretation. We thank Bertrand Ducos for virus titration. **Dudok group**. We thank the Bioengineering Core at Baylor College of Medicine for support with research instrumentation. **Grant support.** The project was supported by the Klingenstein-Simons Fellowship Award in Neuroscience (FSP); the McNair Medical Foundation (FSP, BD); Welch Foundation grant Q-2016-20190330 (FSP), the Whitehall foundation (BD); NIH grants R01EB027145 (FSP), U01NS113294 (FSP, MAL), U01NS118288 (FSP), U01NS133971 (FSP, LB), R00NS117795 (BD), R01NS136027 (FSP, JR), U01NS118300 (NJ), U01NS137449 (NJ), RF1NS128901 (JR), R34NS132045 (JR); NSF NeuroNex grant 1707359 (FSP) and IdeasLab grant 1935265 (FSP).

## Author contributions

**St-Pierre lab.** FSP conceived the project. FSP and MAL coordinated the project. ZL, XL, HL, KLC, and AM maintained and improved the screening platform hardware and data analysis code. MAL modified screening protocols; MAL and MS characterized screening protocols. MAL performed GEVI screening with help from SY and AM. SY conducted confocal microscopy data. XL and MAL conducted all other *in vitro* benchmarking experiments. XD, SZ, MAL and SL cloned screening libraries and viral vectors; MAL and FSP prepared figures and wrote the manuscript. **Bourdieu lab.** AA and JB generated the JEDI-2P AAV. CMH and VV performed mouse surgeries. VV carried out experiments, analyses and figure production. VV, JB, and LB wrote text, captions and discussed critical aspects of the project. **Ji lab.** JZhu prepared samples, acquired data, analyzed the data, and made figures. RGN performed mouse surgeries. J Zhong built the FACED scanning module and wrote the data acquisition program. NJ analyzed the data, made figures and supervised the project. All authors contributed to the writing of the manuscript. **Dudok lab.** BD supervised experiments, analyzed data, prepared figures, and contributed to the manuscript. MM performed experiments, recorded and processed data, prepared figures, and contributed the manuscript. ST performed experiments. **Reimer lab.** JR supervised experiments and analysis, MG, RK, and NH collected optical data, RGL, MG, CLS, and MH performed analyses and generated figures. JR and MG contributed to the manuscript.

