## Supplementary figures for "Designer indicators for two-photon recording of subthreshold voltage dynamics"

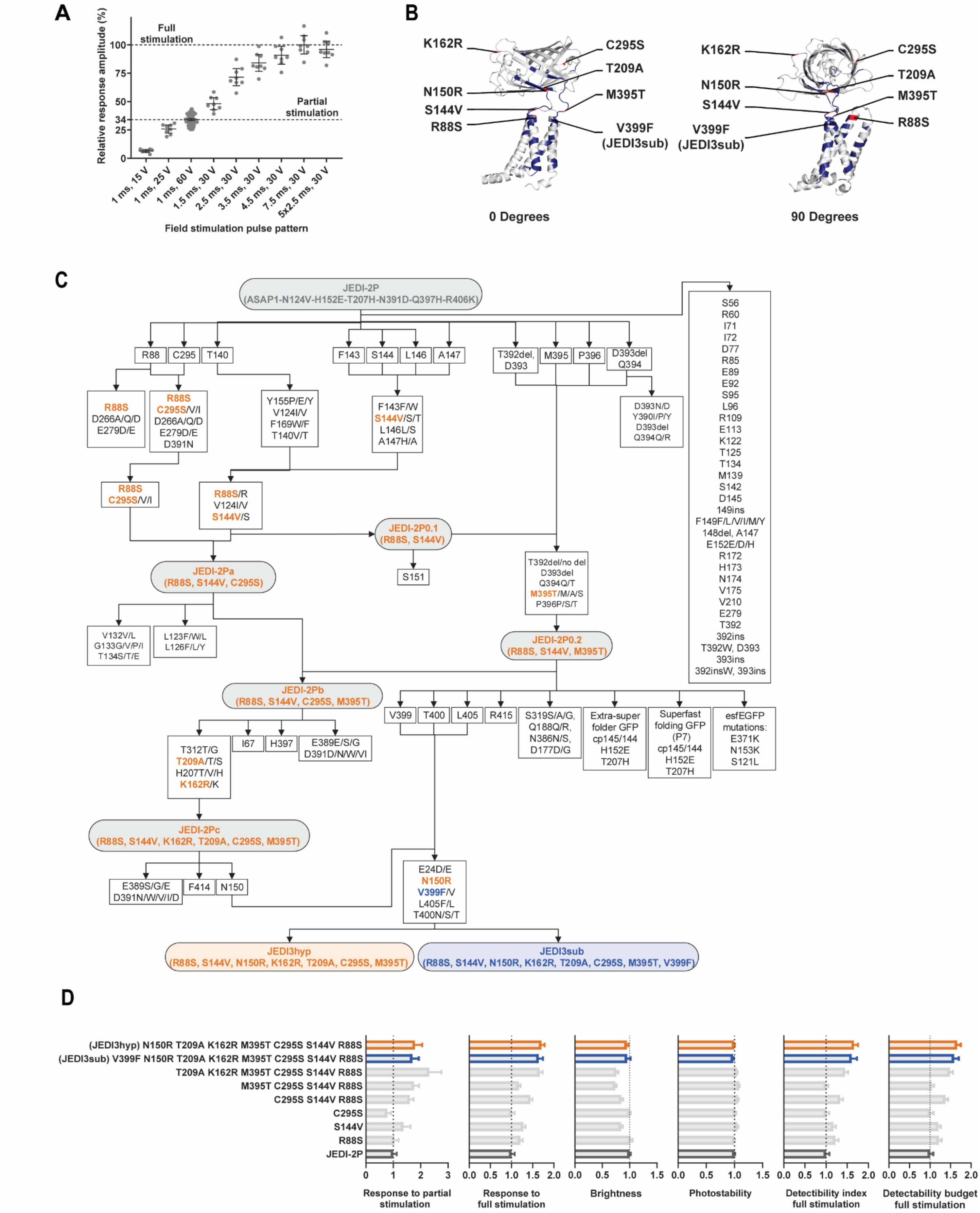
A
Relative response amplitude (%)
Full stimulation
Partial stimulation
100
75
50
34
25
0
1 ms, 15 V
1 ms, 25 V
1 ms, 60 V
1.5 ms, 30 V
2.5 ms, 30 V
3.5 ms, 30 V
4.5 ms, 30 V
7.5 ms, 30 V
5x2.5 ms, 30 V
Field stimulation pulse pattern
B
K162R
C295S
T209A
N150R
M395T
S144V
R88S
V399F (JEDI3sub)
0 Degrees
90 Degrees
C
JEDI-2P (ASAP1-N124V-H152E-T207H-N391D-Q397H-R406K)
R88
C295
T140
F143
S144
L146
A147
T392del, D393
M395
P396
D393del Q394
R88S D266A/Q/D E279D/E
R88S C295S/V/I D266A/Q/D E279D/E D391N
Y155P/E/Y V124I/V F169W/F T140V/T
F143F/W S144V/S/T L146L/S A147H/A
D393N/D Y390I/P/Y D393del Q394Q/R
S56 R60 I71 I72 D77 R85 E89 E92 S95 L96 R109 E113 K122 T125 T134 M139 S142 D145 149ins F149F/L/V/I/M/Y 148del, A147 E152E/D/H R172 H173 N174 V175 V210 E279 T392 392ins T392W, D393 393ins 392insW, 393ins
R88S C295S/V/I
R88S/R V124I/V S144V/S
JEDI-2P0.1 (R88S, S144V)
S151
T392del/no del D393del Q394Q/T M395T/M/A/S P396P/S/T
JEDI-2Pa (R88S, S144V, C295S)
V132V/L G133G/V/P/I T134S/T/E
L123F/W/L L126F/L/Y
JEDI-2P0.2 (R88S, S144V, M395T)
JEDI-2Pb (R88S, S144V, C295S, M395T)
V399
T400
L405
R415
S319S/A/G, Q188Q/R, N386N/S, D177D/G
Extra-super folder GFP cp145/144 H152E T207H
Superfast folding GFP (P7) cp145/144 H152E T207H
esfEGFP mutations: E371K N153K S121L
T312T/G T209A/T/S H207T/V/H K162R/K
I67
H397
E389E/S/G D391D/N/W/V/I
JEDI-2Pc (R88S, S144V, K162R, T209A, C295S, M395T)
E389S/G/E D391N/W/V/I/D
F414
N150
E24D/E N150R V399F/V L405F/L T400N/S/T
JEDI3hyp (R88S, S144V, N150R, K162R, T209A, C295S, M395T)
JEDI3sub (R88S, S144V, N150R, K162R, T209A, C295S, M395T, V399F)
D
(JEDI3hyp) N150R T209A K162R M395T C295S S144V R88S
(JEDI3sub) V399F N150R T209A K162R M395T C295S S144V R88S
T209A K162R M395T C295S S144V R88S
M395T C295S S144V R88S
C295S S144V R88S
C295S
S144V
R88S
JEDI-2P
Response to partial stimulation
Response to full stimulation
Brightness
Photostability
Detectibility index full stimulation
Detectability budget full stimulation

**Figure S1. 2P directed evolution of GEVIs for enhanced responses to subthreshold voltage dynamics.**
(**A**) Changing electrical field stimulation strength and duration depolarize HEK293-Kir2.1 cells to different extents, as visualized by JEDI-2P fluorescence. The partial stimulation pattern was used to screen for enhanced responses to subthresholds, while the full stimulation pattern was used to determine GEVIs' maximal responses. (**B**) Residues targeted during directed evolution, highlighted on an *in-silico* 3D model. Residues mutated in JEDI3 indicators are shown in red, except for V399F, which is only found in JEDI3sub. Sites in blue were screened but did not yield mutations incorporated into the final JEDI3 sensors. (**C**) Directed evolutionary path leading to the identification of JEDI3hyp (orange) and JEDI3sub (all orange plus blue) from the parental sensor, JEDI-2P (gray). Intermediate candidates are shown in orange with a gray background. Some combination libraries included mutations from the literature or screens of other sensor lines. (**D**) Comparative performance of JEDI3sub, JEDI3hyp, and selected intermediates in our 2P multiparametric screen. Values are normalized to JEDI-2P. The detectability index is calculated as the product of response amplitude and the square root of relative brightness. The detectability budget is defined as the detectability index times the square root of the photostability. Data are presented as mean ± 95% CI, with 7–8 independent transfections per variant.

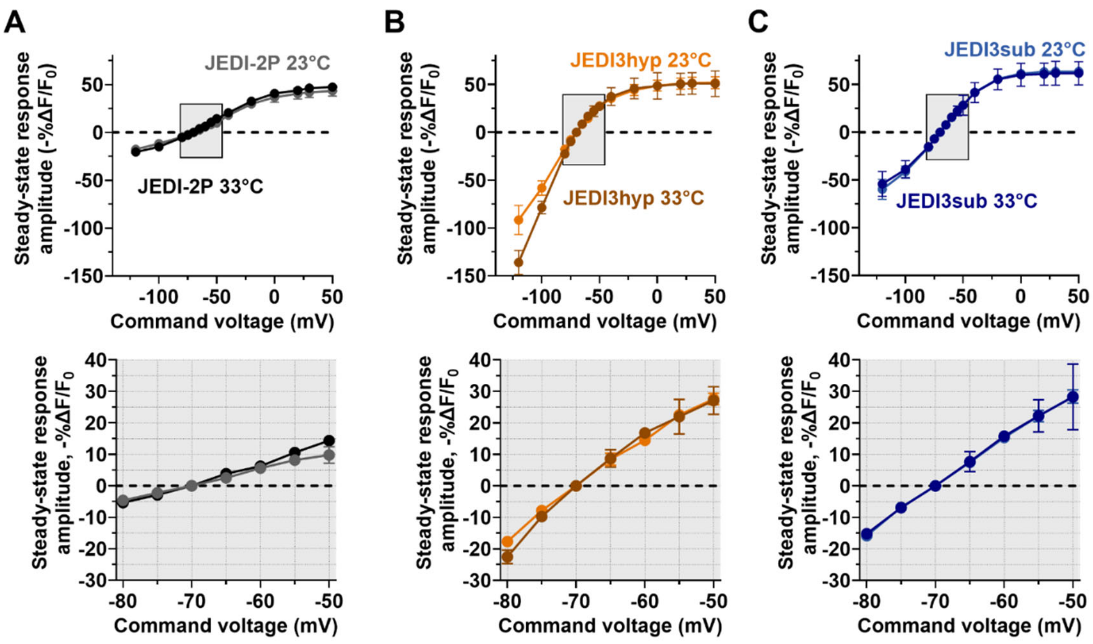


**Figure S2. Effect of temperature on the voltage sensitivity curve.**

Mean fluorescence responses to voltage steps at either 23ºC or 33ºC for (**A**) JEDI-2P, (**B**) JEDI3hyp, and (**C**) JEDI3sub. The bottom row shows additional quantification of smaller voltage steps around subthreshold levels. Imaging was conducted using 2P resonant scanning at a framerate of 440 Hz and setting the laser to 940nm. Voltage clamp was performed in HEK293A cells. 23ºC data: n = 7-10/sensor (same cells as in Fig. 1H). 33ºC data: n = 2 (JEDI-2P) and 3 for JEDI3hyp and JEDI3sub. For all figures, error bars denote 95% CI.

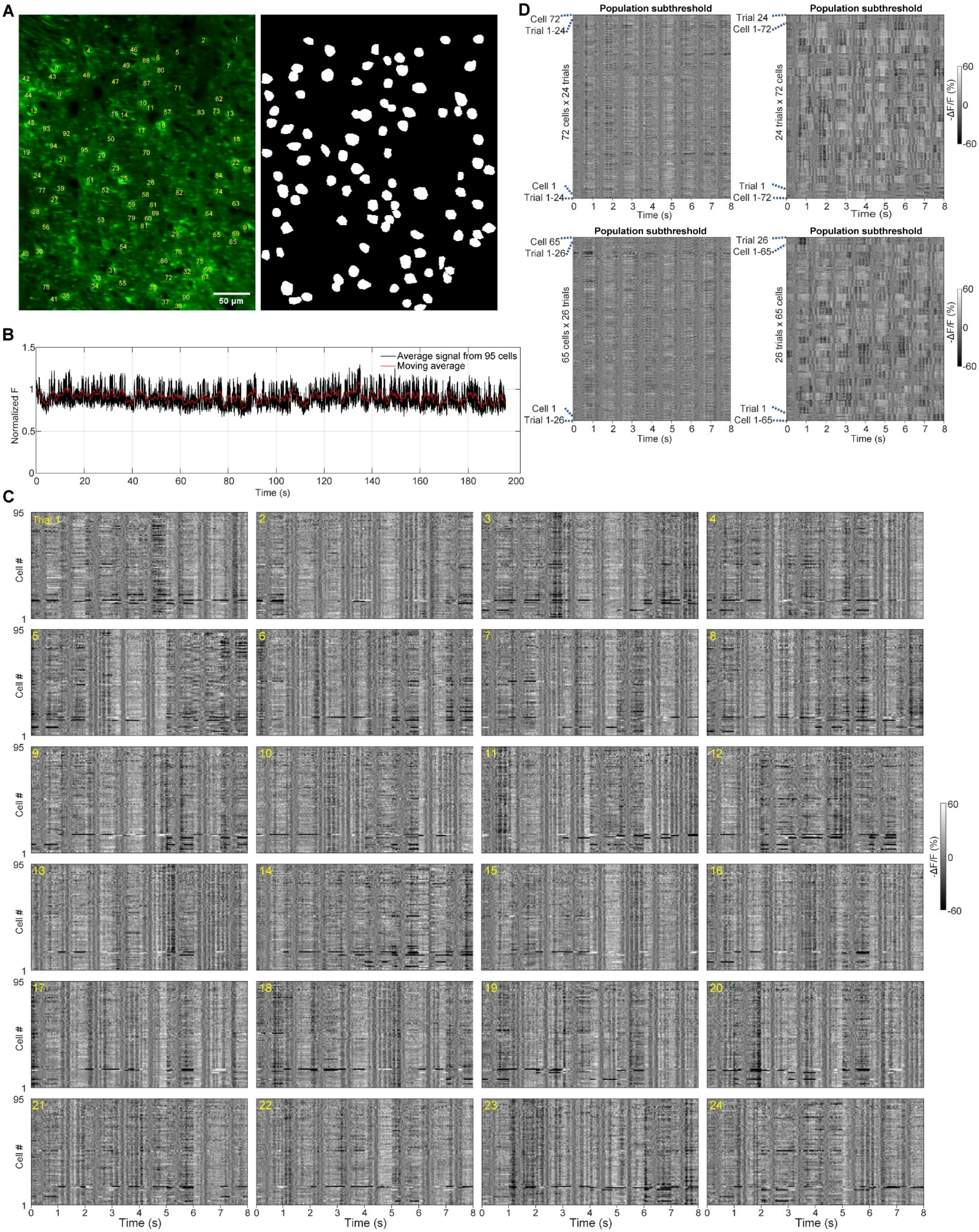
A
50 μm
B
Average signal from 95 cells
Moving average
Normalized F
Time (s)
C
Trial 1
Cell #
Time (s)
-ΔF/F (%)
60
0
-60
D
Population subthreshold
Cell 72
Trial 1-24
72 cells x 24 trials
Cell 1
Trial 1-24
Trial 24
Cell 1-72
24 trials x 72 cells
Trial 1
Cell 1-72
Cell 65
Trial 1-26
65 cells x 26 trials
Cell 1
Trial 1-26
Trial 26
Cell 1-65
26 trials x 65 cells
Trial 1
Cell 1-65
Time (s)
-ΔF/F (%)

**Figure S3: JEDI3sub tracks differences in response to a moving gradient across trials.**
(**A**) *Left:* 2P fluorescence image of L2/3 neurons in a representative field-of-view with regions-of-interest (ROI) indices. *Right*: cell masks. The 2P image is as in Fig. 3B. (**B**) JEDI3sub showed minimal photobleaching over the ~200-s imaging session, as demonstrated by the mean fluorescence signal of 95 cells (black) and its moving average (red). Before averaging, each trace was normalized to its initial fluorescence value. (**C**) Subthreshold response traces from 95 cells across all 24 trials. (**D**) Two additional fields-of-view from separate mice showing single-trial subthreshold traces grouped by cells (*left*) or trial (*right*).

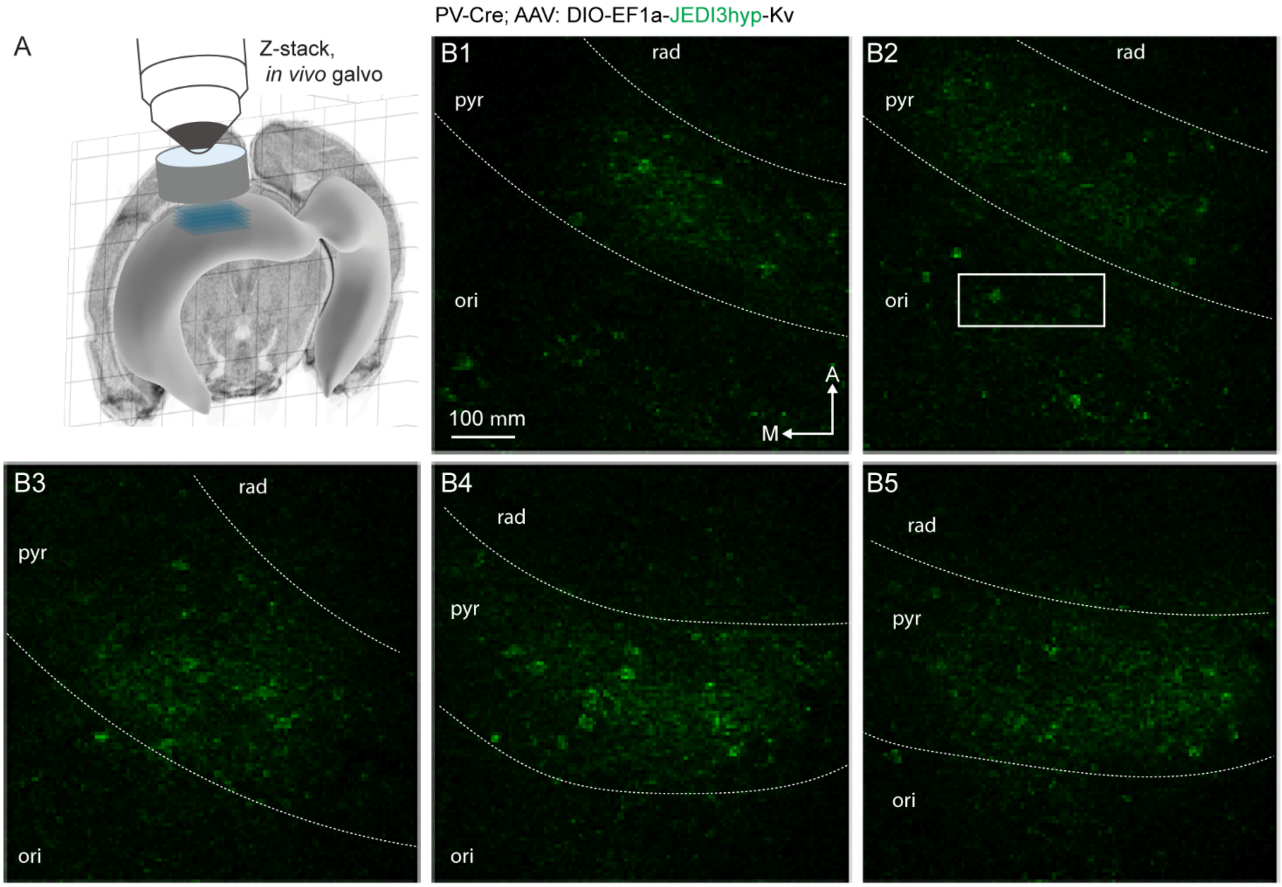


**Figure S4: Representative expression pattern of JEDI3hyp in PV interneurons.**
(**A**) Schematic representation of the hippocampus and the CA1 region where a z-stack of images (blue) was captured. Image source: Allen Mouse Brain Atlas – Brain Explorer 2. (**B1-5**) Cropped regions from example optical sections are shown. Sections were separated by 30 µm in depth (Z). B2's focal plane was adjusted by 10 µm to include the cells (c1-c3) shown in Fig. 4B (solid rectangle). Each slice was cropped at different lateral positions to follow the curvature of the hippocampus and the extent of labeling. Dashed lines show estimated layer borders. Representative of 4 mice. Arrow: M, medial; A, anterior. Layers: ori, oriens; pyr, pyramidal; rad, radiatum.

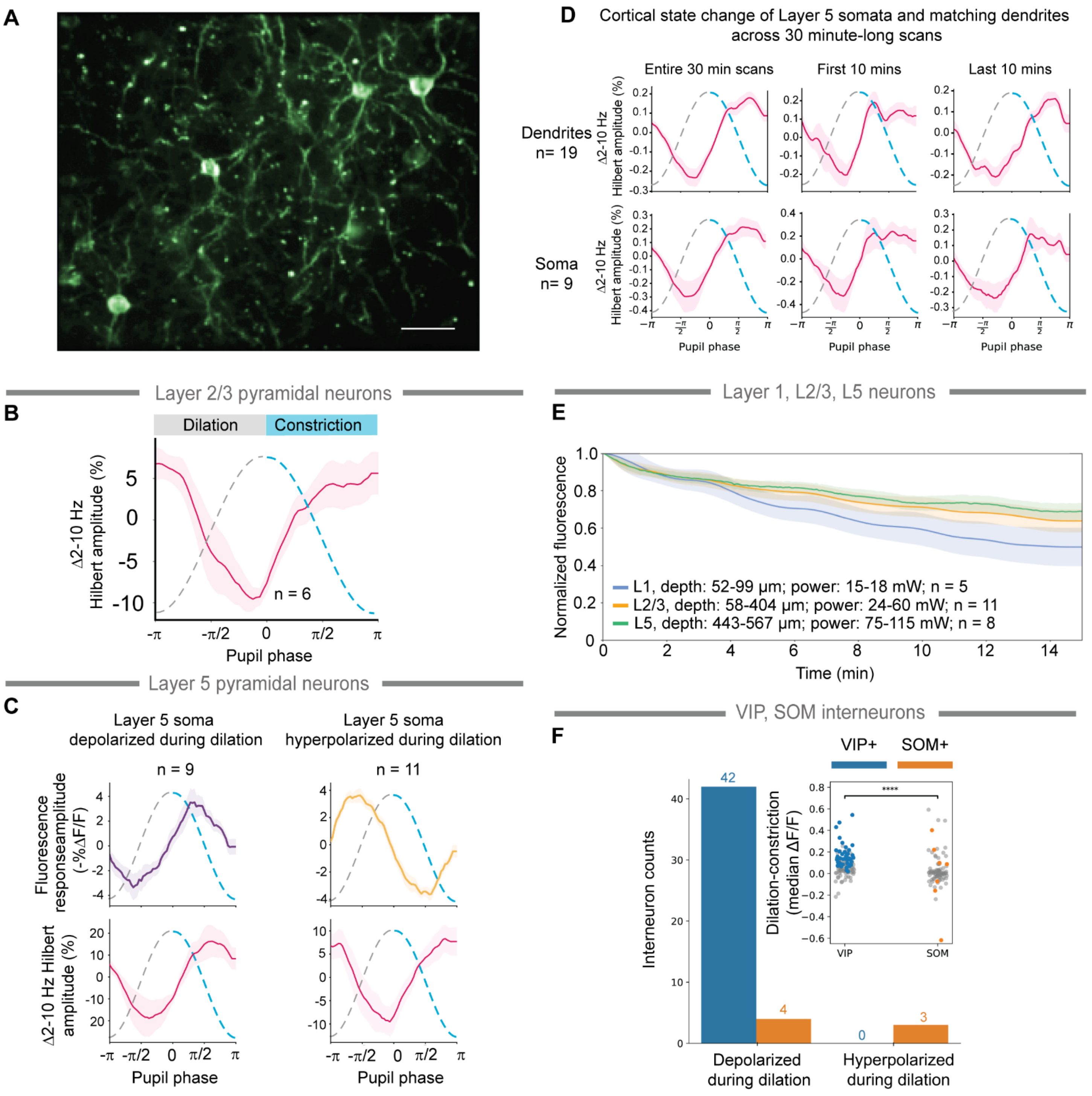


**Figure S5. JEDI3hyp can report subtle subthreshold changes across cell types and cortical layers over extended durations.**

(**A**) Example of JEDI3hyp expression in L5 IT-PN imaged at 87 Hz 434 µm from the surface. Scale bar, 25 um. (**B**) Pupil dilation onset is associated with a reduction in low-frequency activity in Layer 2/3 pyramidal neurons (n = 6), consistent with findings from[1]. Panels B, C, and E were acquired using a 2P resonant scanning microscope. (**C**) Layer 5 pyramidal neurons can be categorized into two functional subclasses based on their response to dilation: hyperpolarized (purple, n = 9) or depolarized (yellow, n = 11). Despite their differing $\Delta F/F_0$ responses to pupil dilation (top panels, Kruskal-Wallis test, $p < 0.05$), both subclasses exhibit similar changes in low-frequency activity (bottom panels, red). (**D**) Changes in low-frequency activity in Layer 5 somas (bottom panel) and corresponding dendrites (top panel) when analyzing the entire 30-minute scans (first column). Fluorescence changes associated with pupil phase remain stable, as shown by similar profiles between the first (center column) and last (third column) 10 minutes of imaging. Images were acquired using a 2P-RAM mesoscope. (**E**) JEDI3hyp demonstrates high photostability over 15 minutes of imaging. Mean values are shown, and shaded areas are the SEM. The power was measured post-objective. (**F**) JEDI3hyp captures interneuron properties previously observed in whole-cell patching data[1]. VIP cells are uniformly depolarized during dilation (blue). SOM interneurons displays variable responses to pupil phase (orange), suggesting heterogeneity in functional cell types. Data for (E) was acquired using 2P random-access scanning via acoustic optic deflectors (Femto3D Atlas). All other data was acquired using resonance scanning.
